# Delivery of loaded MR1 monomer results in efficient ligand exchange to host MR1 and subsequent MR1T cell activation

**DOI:** 10.1101/2022.07.11.499573

**Authors:** Corinna A. Kulicke, Gwendolyn M. Swarbrick, Nicole A. Ladd, Meghan Cansler, Megan Null, Aneta Worley, Chance Lemon, Tania Ahmed, Joshua Bennett, Taylor N. Lust, Chelsea M. Heisler, Megan E. Huber, Jason R. Krawic, Laurisa M. Ankley, Savannah K. McBride, Fikadu G. Tafesse, Andrew J. Olive, William H. Hildebrand, Deborah A. Lewinsohn, Erin J. Adams, David M. Lewinsohn, Melanie J. Harriff

**Affiliations:** Division of Pulmonary, Allergy, and Critical Care Medicine, Oregon Health & Science University, Portland, OR 97239, USA; Division of Infectious Diseases, Department of Pediatrics, Oregon Health & Science University, Portland, OR 97239, USA; Department of Biochemistry and Molecular Biology, University of Chicago, Chicago, IL 60637, USA; VA Portland Health Care System, Portland, OR 97239, USA; Department of Molecular Microbiology and Immunology, Oregon Health & Science University, Portland, OR 97239, USA; Department of Microbiology and Immunology, University of Oklahoma Health Sciences Center, Oklahoma City, OK 73104, USA; Department of Microbiology and Molecular Genetics, College of Osteopathic Medicine, Michigan State University, East Lansing, MI 48824, USA

**Keywords:** MAIT cells, MAIT cell ligands, MR1-restricted T cells, antigen presentation, cellular immune response, mucosal immunology, T-cell

## Abstract

MR1-restricted T cells have been implicated in microbial infections, sterile inflammation, wound healing and cancer. Similar to other antigen presentation molecules, evidence supports multiple, complementary MR1 antigen presentation pathways. To investigate ligand exchange pathways for MR1, we used MR1 monomers and tetramers loaded with 5-(2-oxopropylideneamino)-6-d-ribitylaminouracil (5-OP-RU) to deliver the antigen. Using MR1-deficient cells reconstituted with wild-type MR1 or MR1 molecules that cannot bind 5-OP-RU, we show that presentation of monomer-delivered 5-OP-RU is dependent on cellular MR1 and requires the transfer of ligand from the soluble molecule onto MR1 expressed by the antigen presenting cell. This mode of antigen delivery strengthens the evidence for post-ER ligand exchange pathways for MR1, which could represent an important avenue by which MR1 acquires antigens derived from endocytosed pathogens.

## Introduction

MR1-restricted T (MR1T) cells respond to small molecule antigens presented in the context of MHC I-related protein 1 (MR1) and include mucosal associated invariant T (MAIT) cells, which express the canonical *TRAV1-2* T cell receptor (TCR) alpha chain^1^. They are increasingly being recognized for their role in early responses to microbial infection, but also in non-infectious conditions including a variety of autoimmune disorders, wound healing and cancer (reviewed e.g. in ^1,2^). Despite the heightened interest in these donor-unrestricted T cells, the mechanisms governing the presentation of MR1T cell antigens remain incompletely defined.

Successful antigen presentation requires the antigen presenting molecule to leave the endoplasmic reticulum (ER), bind an antigen, and display the bound antigen at the cell surface for TCR recognition. How these three features are achieved mechanistically, what compartments they take place in, and in what sequence they occur differs substantially between antigen presentation pathways. For major histocompatibility complex (MHC) I/peptide complexes, antigen binding generally precedes, and is indeed required for, ER egress and followed by translocation to the cell surface (reviewed e.g. in ^3^). MHC II and CD1 molecules, by contrast, exit the ER bound to a surrogate ligand which is exchanged for antigenic ligand in the endosomal compartment after internalization from the cell surface (reviewed e.g. in ^4,5^).

For MR1, an MHC I-like mode of antigen acquisition has been described in which more MR1 associates with β_2_-microglobulin and acquires endoglycosidase H resistance upon addition of ligand to the culture medium^6,7^. In line with this, McWilliam, Mak, et al. have used a fluorescently labelled MR1 antigen analogue to directly demonstrate loading of exogenous ligand in the ER^7^. Studies have linked the mechanism for MR1 ER egress to the charge neutralization of a lysine residue within the MR1 ligand binding groove, which has been suggested to result in a conformational change conducive to ER egress^6,7^. The most potent MR1 ligand described to date, 5-OP-RU, forms a Schiff base with this lysine residue, resulting in its neutralization^6,8^.

In addition, recent findings support the hypothesis that there are complementary, non-exclusive presentation pathways for MR1, allowing the molecule to survey diverse intracellular compartments^9–14^. Multiple lines of evidence suggest that ligand exchange can occur on pre-loaded MR1 molecules and that this may take place in an endosomal compartment. Most notably, releasing MR1 from the ER prior to antigen exposure by pre-incubation with the MR1 ligand 6-formylpterin (6-FP) enhances presentation of exogenous antigenic ligand to MR1T cells^10^. In line with this, reduced expression of proteins involved in the trafficking and recycling of endosomal vesicles such as VAMP4, Rab6^9^, syntaxin-4, syntaxin-16, and VAMP2^10^ impairs MR1-mediated antigen presentation. Based on these observations, we and others have proposed ligand exchange in a post-ER compartment as a complementary mechanism for MR1 ligand acquisition^6,10,11,14^. In a mechanism akin to MHC II and CD1 molecules, we hypothesize that MR1 molecules internalized from the cell surface can be re-loaded with microbial antigens in the endosomal compartment and recycle to the cell surface to activate MR1T cells. Here we investigate the mechanisms underlying MR1 ligand exchange using a novel delivery method for MR1T cell antigens. We find that MR1 ligands are efficiently presented when delivered in the context of soluble MR1 molecules. This presentation is dependent on host MR1 and involves the transfer of antigen from the recombinant molecule onto MR1 protein expressed by the cell. Building upon our previously described tetraSPOT assay^15^, this method, the exSPOT, extends the tool box available to investigate MR1 biology and provides important insights into the mechanisms of MR1-mediated antigen presentation.

## Results

### MR1 ligands delivered to antigen presenting cells on soluble MR1 molecules are efficiently recognized by an MR1T cell clone

The MR1 ligand 5-OP-RU is inherently unstable in aqueous environments^8,16^, which presumably includes the intracellular milieu. However, the formation of a covalent bond with the lysine residue at position 43 of MR1 stabilizes the antigen^8^. We have successfully used such MR1/5-OP-RU complexes in their tetramerized form in plate-bound assays to activate MR1T cells^15^. Similarly, Priya and Brutkiewicz have used latex beads coated with MR1/5-OP-RU complexes to expand and activate human MAIT cells^17^. While these studies used recombinant MR1 molecules immobilized on a scaffold, here we sought to improve the efficiency of ligand delivery into physiologically relevant antigen presenting cells (APCs) using soluble MR1.

To test this system, we incubated human dendritic cells (DCs) with MR1 tetramers loaded with the canonical ligand 5-OP-RU (exSPOT), or applied the same concentration of tetramer directly to the ELISPOT plate prior to adding an MR1T cell clone (tetraSPOT). Intriguingly, the MR1T cells readily produced interferon-γ (IFN-γ) in response to the DCs incubated with soluble MR1/5-OP-RU in a dose-dependent manner (Figure 1a). As expected based on previous work^15^, they also produced IFN-γ in the tetraSPOT. A quantitative comparison between the two assays was not possible due to unknown coating efficiency in the tetraSPOT. MR1T cells did not respond to the MR1/6-FP tetramer in the tetraSPOT, but we observed MR1T responses to the MR1/6-FP tetramers incubated with DCs at the highest concentrations (Figure 1a). Similar to the tetraSPOT^15^, MR1T cell activation by DCs incubated with soluble MR1 tetramers was dependent on MR1, as blocking with the anti-MR1 antibody clone 26.5 markedly reduced the response (Figure 1b, EC_50_ of 0.95 pM *vs* 31.62 pM, p<0.0001). Thus, 5-OP-RU delivered in the context of soluble, recombinant MR1 is efficiently presented by DCs and activates a human MR1T cell clone in an MR1-dependent manner.

**Fig 1.**
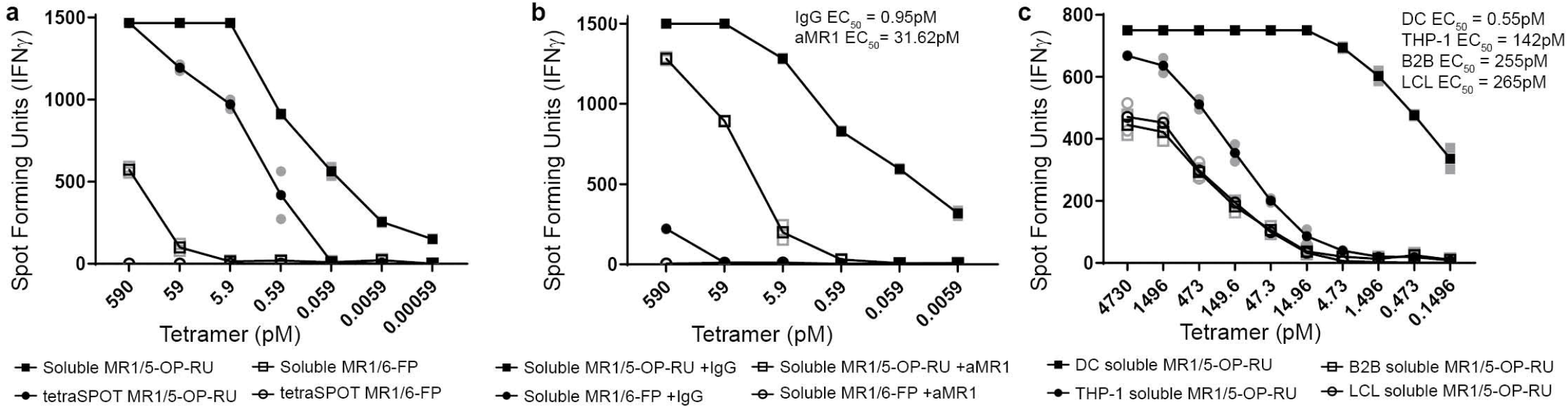
MR1 ligands delivered to antigen presenting cells on soluble MR1 molecules are efficiently recognized by MR1T cell clones. **a.** MR1T cell clone (1e4) IFN-γ responses to MR1/5-OP-RU or MR1/6-FP tetramer, either added to DCs (1e4) (exSPOT) or bound to ELISPOT plate (tetraSPOT). **b.** MR1T cell clone (1e4) IFN-γ response to MR1/5-OP-RU is inhibited by anti-MR1 antibody blockade (aMR1, 2 μg/ml). **c**. MR1T cell clone (1e4) IFN-γ response to MR1/5-OP-RU tetramer, added to either DCs, THP-1 cells, BEAS-2B (B2B) cells or B-LCL (1e4). EC_50_ calculated using non-linear regression and the difference in the EC_50_ was statistically significant between the curves (p<0.0001) by the extra sum-of-squares F test. Data are representative of n=3 independent experiments each. Technical duplicates within the representative experiment are shown with a line connecting the mean. For EC_50_ values of replicate experiments see Supplementary Tables 1+2.

As professional APCs, DCs are particularly well equipped for phagocytosis of particulates of various sizes and cross-presentation^18^. To investigate whether such specific functionality was required in the exSPOT, we incubated the monocytic cell line THP-1, a B cell line (LCL), or the airway epithelial cell line BEAS-2B with MR1/5-OP-RU tetramers and quantified IFN-γ responses. As with DCs, the MR1T cell clone produced IFN-γ in response to all of the tetramer-incubated cell types (Figure 1c). However, DCs were nearly three orders of magnitude more efficient at presenting the antigen compared to the other cell types (EC_50_ of 0.55 pM *vs* 142 – 265 pM, p<0.0001).

### Ligand delivered in the context of MR1 monomer is presented more efficiently than free ligand

Next, we compared the presentation of MR1/5-OP-RU tetramers and MR1/5-OP-RU monomers to reduce the variability associated with tetramerization (e.g. biotinylation efficiency, streptavidin binding, and ligand saturation). Using the MR1 monomer further circumvented potential variability introduced by instability of the tetramer complex over time.

To our surprise, presentation of ligand from monomer-incubated DCs to MR1T cells was even more efficient than presentation of ligand from tetramer-incubated DCs, when accounting for valency (Figure 2a). Although the EC_50_ values of the complexes were comparable (Tetramer EC_50_: 0.204 pM *vs* Monomer EC_50_: 0.22 pM, p=0.7517), there was a marked difference in the estimated EC_50_ for 5-OP-RU which takes into account that each tetramer carries up to four 5-OP-RU molecules whereas each monomer carries one at most (Tetramer 5-OP-RU EC_50_: 0.816 pM *vs* Monomer 5-OP-RU EC_50_: 0.22 pM, p<0.0001). Comparing monomer-delivered 5-OP-RU to free ligand further revealed improved efficiency of the monomer over the unbound antigen (Figure 2a; Monomer EC_50_: 0.22 pM *vs* Free EC_50_: 4.97 pM, p<0.0001). Of note, these calculations assume full occupancy of monomers and tetramers.

**Fig 2.**
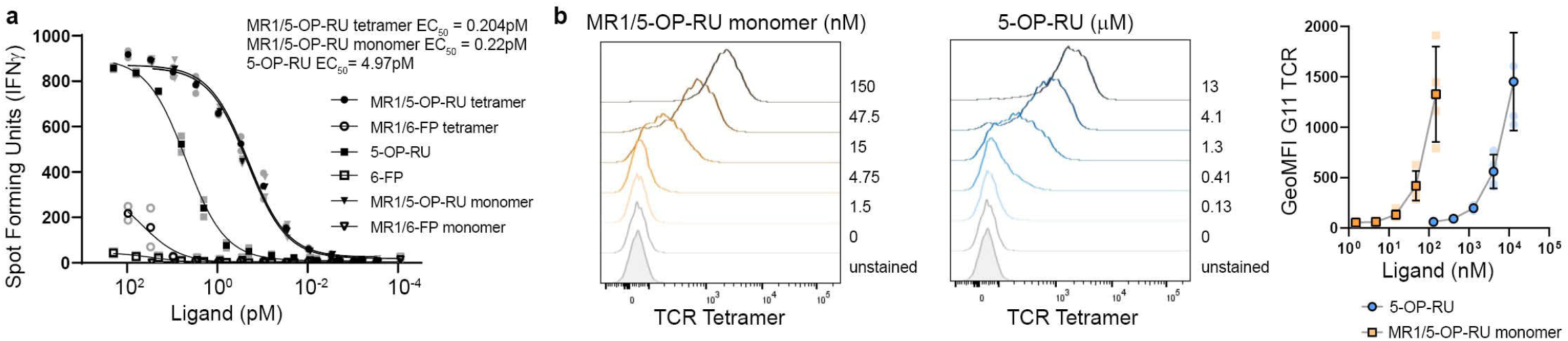
Ligand delivered in the context of MR1 monomer is presented more efficiently than free ligand. **a.** MR1T cell clone (1e4) IFN-γ response to DCs incubated with MR1/5-OP-RU tetramer, MR1/5-OP-RU monomer, and soluble 5-OP-RU. EC_50_ calculated using non-linear regression. The difference in the EC_50_ was statistically significant between the MR1/5-OP-RU tetramer and 5-OP-RU and between MR1/5-OP-RU monomer and 5-OP-RU curves (p<0.0001), but there was no significant difference in the EC_50_ of MR1/5-OP-RU tetramer and monomer by the extra sum-of-squares F test (p=0.7486). Data are representative of n=3 independent experiments. Technical duplicates within the representative experiment are shown with a line depicting the non-linear regression curve. For EC_50_ values of replicate experiments see Supplementary Table 3. **b.** BEAS-2B.MR1-GFP cells were incubated with the indicated concentrations of MR1/5-OP-RU monomer or free 5-OP-RU overnight and surface stained for flow cytometry with APC-conjugated TCR tetramer. Representative histograms are shown with pooled geometric mean fluorescence intensity (GeoMFI) from n=4 experiments. The same unstained and untreated control are shown in both histogram plots for easy comparison. Error bars denote SD of the mean of pooled experiments which are shown individually in lighter colors.

To further investigate the relative efficiency of loading free *vs* monomer-bound 5-OP-RU, we wanted to specifically detect MR1 molecules bearing 5-OP-RU at the cell surface by flow cytometry. To this end, we generated a TCR tetramer based on the TCR sequence of the same previously characterized MR1T cell clone that was used in the functional assays (*TRAV1-2*^+^ clone D426-G11 – see references^15,19^). We confirmed that this reagent specifically stained MR1 molecules loaded with 5-OP-RU but not 6-FP (Supplementary Figure 1). When we incubated BEAS-2B cells overexpressing green fluorescent protein (GFP)-tagged MR1 (BEAS-2B.MR1-GFP) with either MR1/5-OP-RU monomer or free 5-OP-RU overnight and stained with the TCR tetramer to track the cell surface translocation of MR1 bound to the antigen, almost 100-fold higher concentrations of free 5-OP-RU were needed to reach cell surface levels comparable to monomer-delivered ligand (Figure 2b). Together, these results demonstrate that soluble, recombinant MR1 is an efficient mechanism for delivery of MR1T cell antigens in a variety of cell lines *in vitro*.

### Presentation of MR1 ligands delivered on soluble MR1 molecules requires cellular MR1

In principle, the MR1/5-OP-RU complex that interacts with the TCR in our assay could be either endogenously expressed MR1 bound to 5-OP-RU from the monomer or the recombinant MR1/5-OP-RU monomer itself. Thus, we tested whether endogenous MR1 was required. We incubated wild-type (WT) BEAS-2B cells, *MR1*^-/-^ BEAS-2B cells lacking all isoforms of MR1, and *MR1^-/-^* BEAS-2B cells reconstituted with tetracycline-inducible *MR1A*^20,21^ with MR1/5-OP-RU monomer and used them as APCs. As expected, we detected responses to WT BEAS-2B cells incubated with MR1/5-OP-RU monomers (Figure 3a (i)), but when these monomers were incubated with the *MR1^-/-^* BEAS-2B cell line, the response was completely abrogated (Figure 3a (ii)). Reconstitution of MR1 expression in MR1-deficient cells resulted in complete recovery of MR1T cell responses to the monomer-delivered ligand (Figure 3a (iii)). These observations were confirmed in THP-1 cells (Figure 3b).

**Fig 3.**
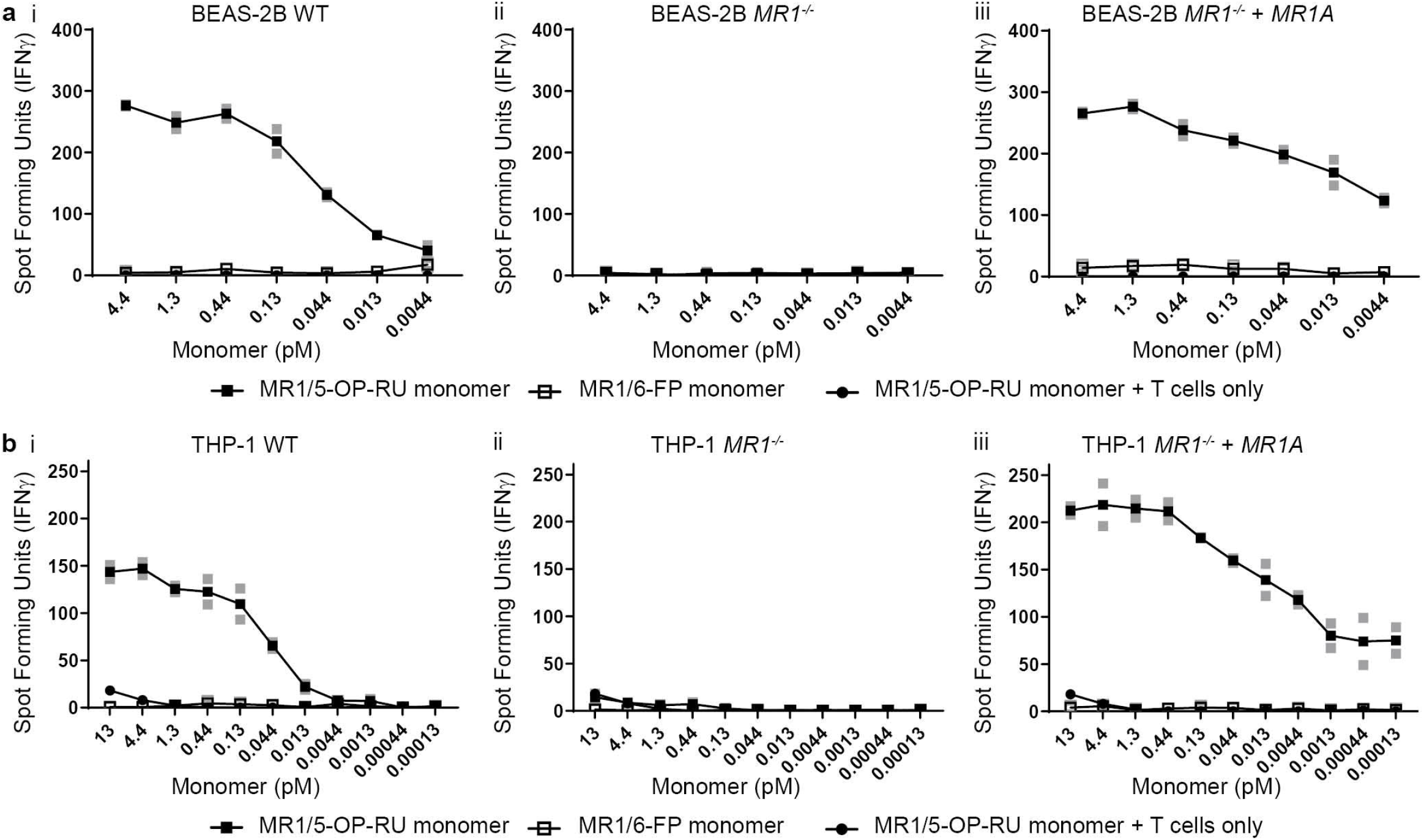
Presentation of MR1 ligands delivered on soluble MR1 molecules requires cellular MR1. **a.** MR1T cell clone (1e3) IFN-γ response to BEAS-2B, BEAS-2B *MR1^-/-^* and BEAS-2B *MR1^-/-^* : *MR1A* (1e4) incubated with MR1/5-OP-RU or MR1/6-FP monomers. Also included MR1T clone IFN-γ response to MR1/5-OP-RU monomer only with no antigen presenting cells. **b.** Same as **a.**, except the antigen presenting cells are THP-1, THP-1 *MR1^-/-^* and THP-1 *MR1^-/-^* : *MR1A* (1e4). Data are representative of n=2 independent experiments. Technical duplicates within the representative experiment are shown with a line connecting the mean.

### Ligands are transferred from soluble MR1 molecules onto cellular MR1

Having established that MR1 expression by the APC itself was required for presentation of monomer-delivered ligand, we considered two possible scenarios: (1) dimerization or indirect binding of the monomer to host MR1 at the cell surface and (2) uptake of the MR1/5-OP-RU complex and transfer of the antigen from the monomer onto cellular MR1.

Since the MR1 molecules used to generate the ligand-loaded monomers and tetramers do not contain a transmembrane domain^8^, their presence on the cell surface and, hence, ability to activate MR1T cells, would require lateral interactions with other membrane-bound proteins. Of note, Lion and colleagues have demonstrated that a GFP-tagged version of the MR1B isoform weakly associated with the MR1A isoform although that observation was considered a potential technical artifact^22^. To directly investigate the presence of the monomer on the cell surface, we incubated BEAS-2B.MR1-GFP cells with the biotinylated MR1/5-OP-RU monomer and stained the cells with fluorophore-coupled streptavidin. We did not detect any signal from the biotinylated monomer although our controls confirmed that we added sufficient monomer to induce MR1 surface translocation and sufficient streptavidin to detect biotinylated molecules at the cell surface (Figure 4a). Interestingly, we detected a small but notable shift in total MR1 surface levels after incubation with the monomer (Figure 4a (iii)). This suggests that enough 5-OP-RU ligand is delivered to increase not just TCR-reactive, but overall MR1 surface levels. Together, these data do not support MR1 dimer formation at the cell surface.

**Fig 4.**
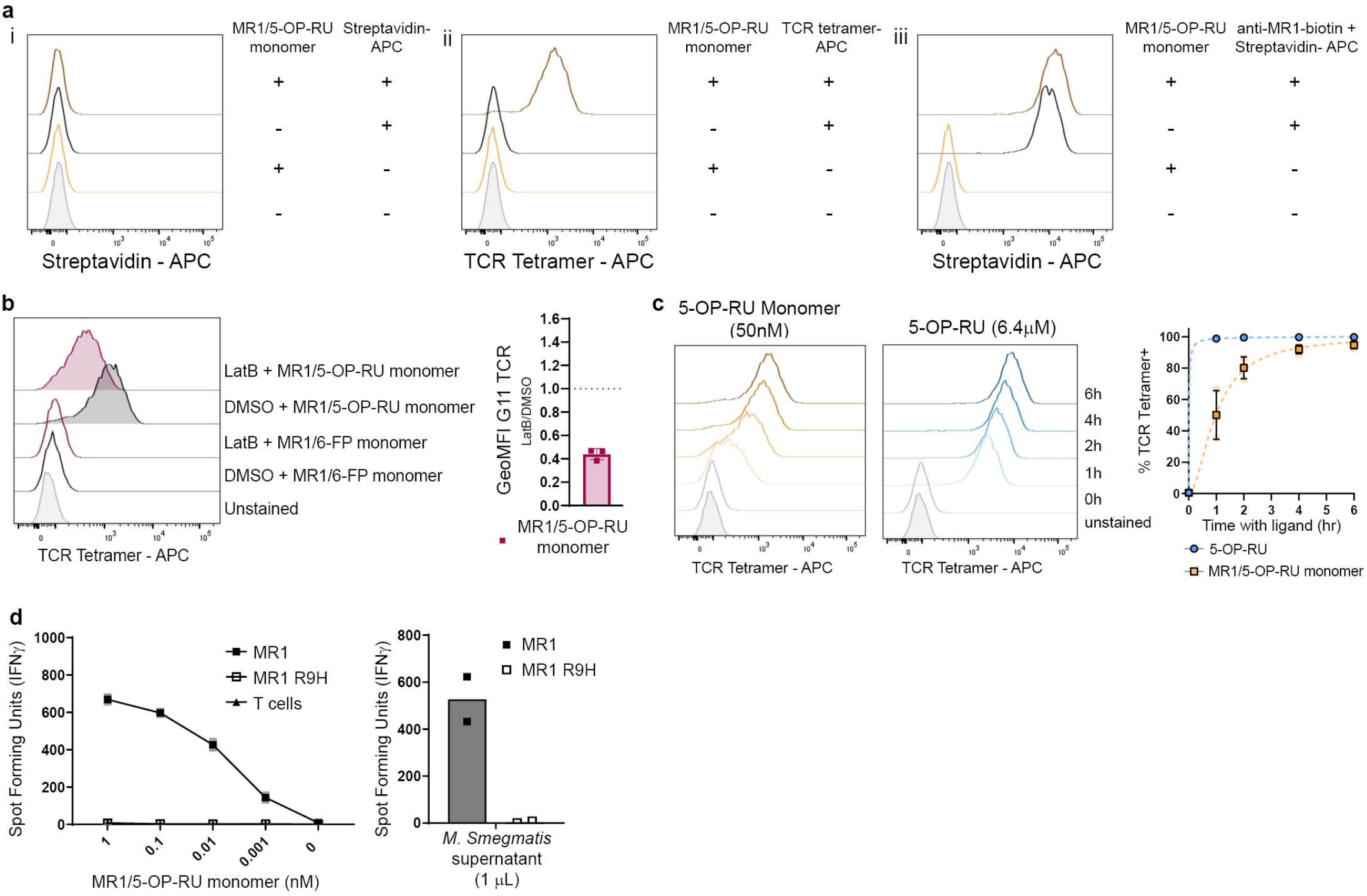
Ligands are transferred from the soluble MR1 molecules onto cellular MR1. **a.** BEAS-2B.MR1-GFP cells (3e5) were pre-incubated with 50 nM of MR1/5-OP-RU monomer overnight before staining with either Streptavidin (i), TCR tetramer (ii) or biotinylated 26.5 anti-MR1 antibody followed by Streptavidin (iii). The same unstained sample is shown in all panels and the same primary antibody only controls are shown in (i) and (iii) for easy comparison. Data are representative of n=3 independent experiments. **b.** BEAS-2B.MR1-GFP cells were pre-incubated with 10 μM of latrunculin B (LatB) for 1 h before addition of MR1/5-OP-RU or MR1/6-FP monomer for an additional 4 h. Cells were surface stained with TCR tetramer. Representative histograms are shown with pooled geometric mean fluorescence intensity (GeoMFI) from n=3 experiments. The same unstained and untreated control are shown in both histogram plots for easy comparison. Error bars denote SD. Bar denotes mean. **c**. BEAS-2B.MR1-GFP cells were incubated with either 50 nM MR1/5-OP-RU monomer or 6.4 μM of free 5-OP-RU for indicated amounts of time before surface staining for flow cytometry with TCR tetramer. Representative histograms are shown with pooled geometric mean fluorescence intensity (GeoMFI) from n=3 experiments. Error bars denote SD of the mean of pooled experiments which are shown individually in lighter colors. **d.** MR1T cell clone (1e3) IFN-γ response to BEAS-2B *MR1^-/-^* cells (1e4) that had been transfected with the pCI AscI MR1 res or pCI AscI MR1 R9H res plasmid, followed by addition of the MR1/5-OP-RU monomer, or 1 μL of *M. smegmatis* supernatant. Data are representative of n=3 independent experiments. Technical duplicates within the representative experiment are shown with a line connecting the mean.

Since we could not find evidence for direct presentation by the monomer, we next tested the hypothesis that the 5-OP-RU is transferred from the soluble MR1/5-OP-RU complex onto the MR1 expressed by the APC. First, we sought to establish whether actin-dependent internalization of soluble MR1 molecules was required for the presentation of monomer-delivered 5-OP-RU. Thus, we treated BEAS-2B.MR1-GFP cells with latrunculin B (LatB), a well-characterized inhibitor of actin polymerization. Surface levels of 5-OP-RU-loaded MR1 were reduced in cells treated with LatB as compared to the dimethyl sulfoxide (DMSO) control (Figure 4b). Of note, LatB treatment also reduced surface expression of TCR-reactive MR1 in response to free 5-OP-RU (Supplementary Figure 2a). Total MR1 as measured by the GFP expression was not reduced by LatB treatment (Supplementary Figure 2b). We concluded that an active actin cytoskeleton is required for presentation of the monomer-derived ligand. This is consistent with previous results demonstrating a requirement for actin for an efficient MR1 response to *Escherichia coli (E.coli)*^13,23^ and could indicate a requirement for either internalization or intracellular trafficking.

Additional evidence that intracellular processing is required came from the observation that the kinetics of presentation differ between monomer-delivered ligand compared to free 5-OP-RU. Here, we incubated BEAS-2B.MR1-GFP cells with concentrations of MR1/5-OP-RU monomer (50 nM) and free 5-OP-RU (6.4 μM) that resulted in intermediate surface expression (Figure 2b) and followed MR1 surface translocation over time. In agreement with McWilliam et al., who detected surface expression of 5-OP-RU-loaded MR1 after 2 h^6^, we saw loaded MR1 as soon as 1 h after addition of the free ligand (Figure 4c). By contrast, it took 6 h for the MR1 surface expression to reach comparable levels after addition of the MR1/5-OP-RU monomer (Figure 4c). These kinetics are consistent with a longer time needed for internalization, processing, ligand exchange, and recycling of monomer-delivered ligand compared to free 5-OP-RU.

To directly investigate transfer onto cellular MR1, we tested whether the MR1 expressed by the APC needed to be capable of presenting 5-OP-RU or whether its presence was sufficient. We reconstituted the *MR1^-/-^* BEAS-2B cell line with a CRISPR-resistant version of *MR1A* that carried the R9H mutation described by Howson et al.^24^. This mutation allows stable MR1 protein expression and binding of 6-FP but renders the protein incapable of presenting 5-OP-RU (^24^ and Supplementary Figures 3 and 4). Indeed, reconstituting the MR1-deficient cells with this mutant did not rescue the presentation of either monomer-delivered 5-OP-RU or *Mycobacterium smegmatis (M. smegmatis)* supernatant (Figure 4d), further supporting the idea that it is the cellular MR1 that presents the 5-OP-RU ligand.

### Ligand exchange occurs in post-ER compartments and does not require acidification

Our data demonstrate the transfer of monomer-delivered 5-OP-RU onto MR1 expressed by the APC but the mechanism of this exchange is unclear. Our lab has previously shown that pre-incubation with the non-stimulatory ligand 6-FP can increase MR1-mediated antigen presentation, presumably by increasing the pool of MR1 molecules available for ligand exchange^10^. When we applied this concept to the monomer delivery assay, we found a similar effect: releasing MR1 from the ER by pre-incubation with 6-FP decreased the EC_50_ for presentation of monomer-delivered ligand from 1.027 pM to 0.1365 pM (p<0.0001; Figure 5a). Thus, MR1 molecules pre-loaded with a non-stimulating ligand appear to contribute to the activation of MR1T cells after delivery of antigenic ligand on monomers, supporting the notion of ligand exchange. In this context, it is worth noting that 6-FP concentrations multiple orders of magnitude higher were required to inhibit 5-OP-RU presentation in a functional assay^25^, which suggests stronger binding of 5-OP-RU^6^ and may help push the equilibrium towards binding of the activating ligand.

**Fig 5.**
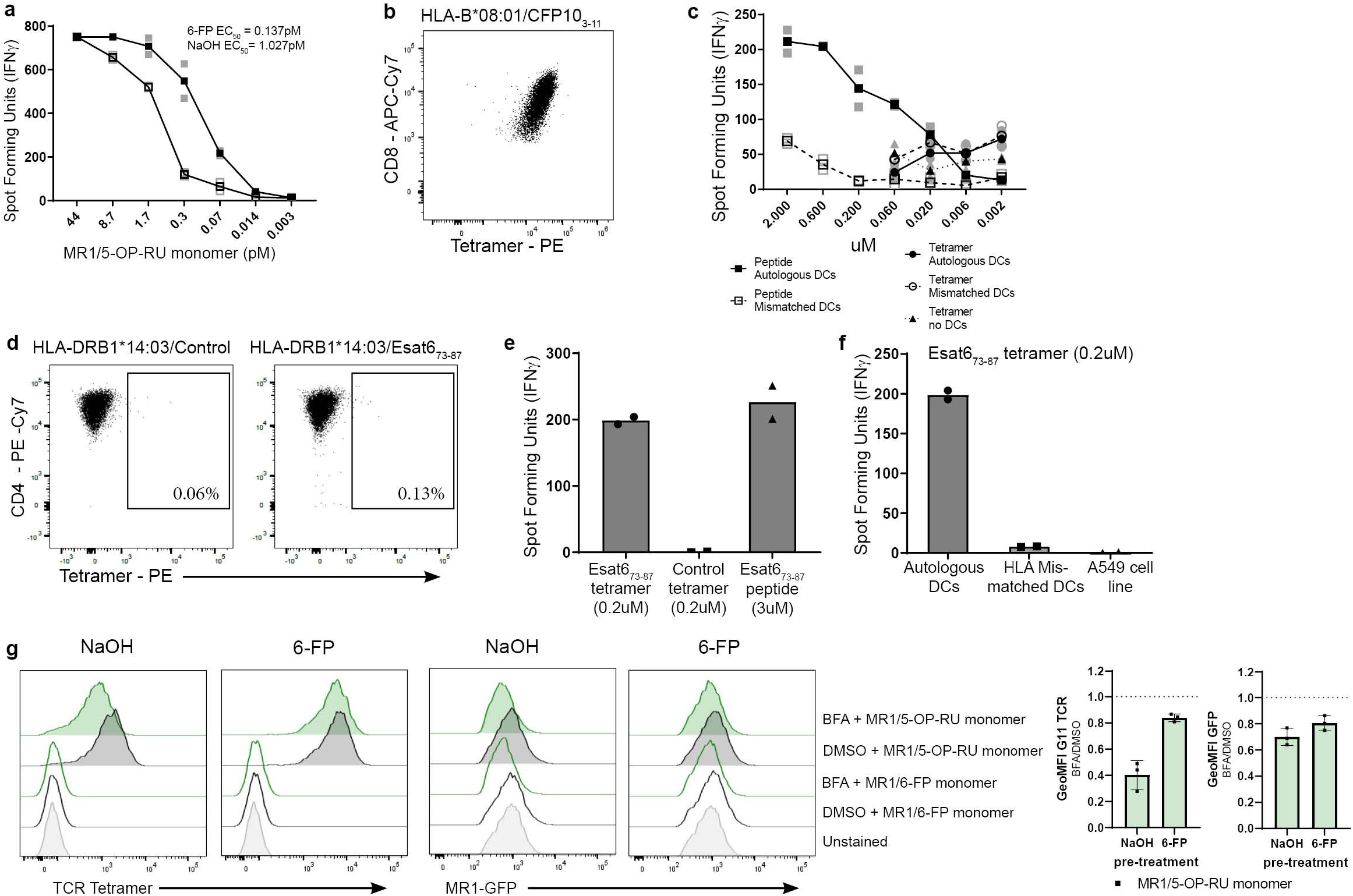
Ligand exchange can occur in post-ER compartments. **a.** MR1T cell clone (1e4) IFN-γ response to BEAS-2B.MR1-GFP (1e4) that were pretreated with 6-FP *vs* NaOH using MR1/5-OP-RU monomer as the antigen. EC_50_ calculated using non-linear regression and the difference in the EC_50_ was statistically significant between the curves (p<0.0001) by the extra sum-of-squares F test. For EC_50_ values of replicate experiments see Supplementary Table 4. **b.** HLA*B08:01-restricted T cell clone D480-F6 was stained with the HLA-B*08:01 tetramer to CFP10_3-11_, along with antibodies to CD3, CD4, CD8 and a live/dead discriminator. Live, CD3^+^CD4^-^CD8^+^ T cells are shown. **c.** HLA*B08:01-restricted T cell clone D480-F6 (5e3) IFN-γ response to DCs (1e4) that were incubated with either the CFP10_3-11_ tetramer or CFP10_3-11_ peptide. 9.5e4 irrelevant T cells (MR1-restricted D426-G11) were also added to the well to decrease T:T presentation of the classical T cell clone. **d.** MHC II-restricted T cell clone D481-D1 was stained with the HLA-DRB1*14:03 tetramer to Esat6_73-87_ or a control tetramer along with antibodies to CD3, CD4, CD8 and a live/dead discriminator. Live, CD3^+^CD8^-^CD4^+^ T cells are shown. **e.** MHC II-restricted T cell clone D481-D1 (2e4) IFN-γ response to D481 (autologous) dendritic cells (2e4) that were incubated with either the Esat6_73-87_ tetramer, control tetramer or Esat6_73-87_ peptide. **f.** MHC II-restricted D481-D1 T cell clone (2e4) IFN-γ response to D481 (autologous) DCs, D520 mismatched dendritic cells or A549 cells (2e4) that were incubated with the Esat6_73-87_ tetramer. **g.** BEAS-2B.MR1-GFP cells were pre-incubated with 6-FP or NaOH overnight. 6-FP was washed off and cells were incubated with 50 ug/ml brefeldin A (BFA) for 1h before addition of MR1/5-OP-RU or MR1/6-FP monomer for an additional 4 h. Cells were surface stained for flow cytometry with TCR tetramer. Representative histograms are shown with pooled geometric mean fluorescence intensity (GeoMFI) from n=3 experiments. The same unstained control is shown in all plots. Error bars denote SD. Bars denote mean. Data in **a**, **c**, **e** and **f** are representative of n=3 independent experiments. In **a** and **c**, technical duplicates within the representative experiment are shown with a line connecting the mean. In **e** and **f**, technical duplicates witin the representative experiment are shown with the bar showing the mean.

To probe to what extent the monomer delivery could be extended to other antigen presentation pathways and potentially gain mechanistic insights from any parallels, we repeated the assay with peptide-loaded MHC I and II tetramers. We first used a fluorophore-conjugated HLA- B*08:01/CFP10_3-11_ tetramer to stain a CD8^+^ T cell clone restricted by this MHC I/peptide complex (D480-F6 – see references ^26^ and ^27^) for flow cytometry, confirming the T cell clone reacted with the tetramer (Figure 5b). However, using the same tetramer to load autologous DCs in an ELISPOT with the same T cell clone did not result in the production of IFN-γ in an APC-dependent manner (Figure 5c). As expected, the T cells responded to DCs incubated with free peptide (Figure 5c). Next, we tested an HLA-DRB1*14:03/ESAT6_73-78_ tetramer in combination with a T cell clone that was specific to the antigen but restricted by a different MHC II allele (D481-D1, from an HLA- DRB1*14:03 negative individual). As expected, this tetramer did not stain the T cell clone by flow cytometry (Figure 5d). Nevertheless, feeding the non-matched tetramer to DCs autologous to the same MHC II-restricted T cell clone elicited an IFN-γ response (Figure 5e). The only source of matched MHC II molecules in this case are the DCs, further supporting a requirement for transfer of the antigen from the soluble protein onto antigen presenting molecules expressed by the APC itself. In line with this, the T cells did not respond when using HLA-mismatched DCs or the A549 cell line which express MHC II alleles different from Donor 481 (^28^, Cellosaurus cell line A-549 (CVCL_0023) (expasy.org)) (Figure 5f). Thus, like with MR1, presentation of the MHC II antigen delivered in the context of recombinant antigen presenting molecules requires transfer onto proteins expressed by the APC.

Since soluble proteins could efficiently deliver peptide for MHC II presentation, we reasoned that the MR1 loading in our assay was similar to the MHC II pathway, which involves exchange of a surrogate ligand in an endosomal compartment (reviewed e.g. in ^4^). To test whether MR1 ligand transfer from the monomer could be mediated in an analogous fashion, we investigated the effect of brefeldin A (BFA) on the presentation of monomer-delivered 5-OP-RU. BFA disrupts the Golgi apparatus and effectively traps secretory proteins in the ER. Indeed, incubation with BFA diminished surface translocation of 5-OP-RU-loaded MR1 in response to the MR1/5-OP-RU monomer (Figure 5g). However, this experiment is unable to distinguish between two different scenarios: 1) ligand exchange occurs in the ER and MR1 needs to subsequently traverse the Golgi for surface presentation, or 2) MR1 needs to leave the ER to be available for exchange in a post-ER compartment. To distinguish between these two hypotheses, we pre-incubated the cells with 6-FP to induce ER egress of a pool of MR1 molecules before adding the BFA. Under these conditions, BFA had a minimal effect on presentation of monomer-delivered ligand (Figure 5g). As expected, presentation of free 5-OP-RU was similarly impaired in the presence of BFA and improved after pre-incubation with 6-FP (Supplementary Figure 5). It is important to note that BFA treatment also reduced total MR1 levels as measured by GFP expression (Figure 5g). These data clearly show that 5-OP-RU delivered on soluble, recombinant MR1 molecules can be transferred to cellular MR1 in a post-ER compartment.

To further characterize the compartment where ligand exchange in the exSPOT occurs, we performed immunofluorescence microscopy. We observed AF647-labeled MR1/5-OP-RU tetramer in compartments labeled with Rab5 (23.51%, standard deviation (SD) = 11.93), an early endosome marker, or LAMP1 (33.1%, SD=12.3), which localizes to late endosomes and lysosomes in these cells (Figure 6a). These observations support the hypothesis that ligand transfer occurs in post-ER compartments, including the early and late endosome.

**Fig. 6.**
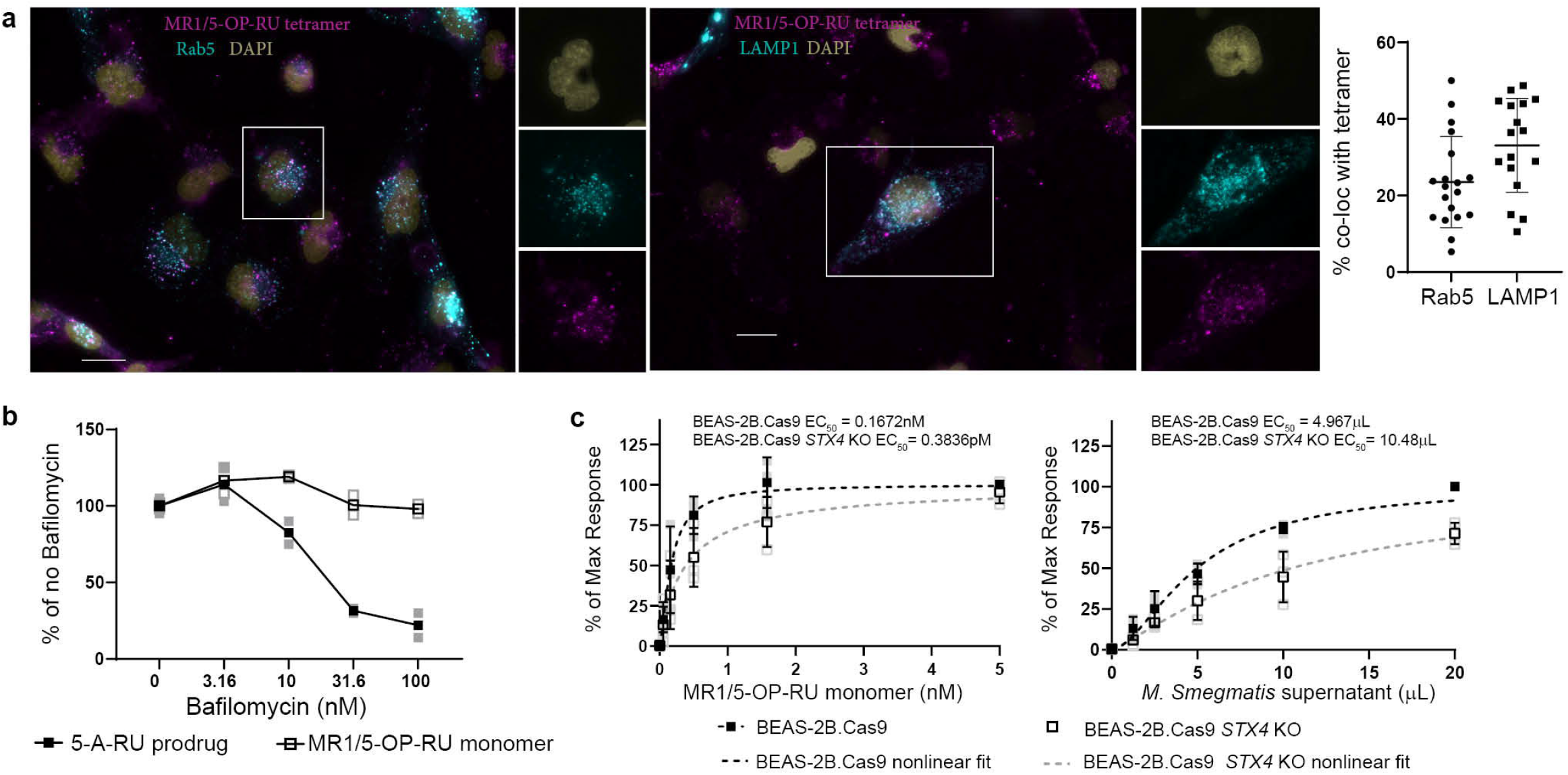
Ligand exchange does not require acidification. **a.** BEAS-2B cells (1e5 or 1e4) were seeded on chamber slides and CellLight RFP reagents for Rab5 and LAMP1 were added and incubated for 18 h at 37C. Tetramer was then added for 2 h at room temperature, followed by washing 2X in PBS. All images acquired were using the same microscope settings. Data are representative of n=3 independent experiments. Error bars denote SD of n=19 cells for Rab5 and n=17 cells for LAMP1 from a representative experiment. Scale bar denotes 20 μm. **b.** MR1T cell clone (5e3) IFN-γ response to BEAS-2B (1e4) pre-treated with Bafilomycin A1 for 2 h, followed by addition of the MR1/5-OP-RU monomer or 5-OP-RU prodrug. Data are representative of n=3 independent experiments. Technical duplicates within the representative experiment are shown with a line connecting the mean. **c**. MR1T cell clone (1e3) IFN-γ response to BEAS-2B.Cas9 or BEAS-2B.Cas9 *STX4* KO cells (1e4) in the presence of the indicated concentrations of *M. smegmatis* supernatant or MR1/5-OP-RU monomer. Data were normalized to the maximum response and pooled from n=3 independent experiments. Error bars denote SD of the mean of pooled experiments which are shown individually in lighter colors. EC_50_ calculated using non-linear regression and the difference in the EC_50_ was statistically significant between the curves (p=0.0032) by the extra sum-of-squares F test.

Exchange of exogenous ligand onto MHC II^4^ and CD1^5^ molecules is potentiated by the low pH in the endolysosomal compartment. Pertinently, endosomal acidification is required for efficient presentation of *E. coli* by MR1 ^13,29^ and McWilliam et al. have proposed that the Schiff base that links 5-OP-RU to the MR1 ligand binding groove is labile under acidic conditions^6,14^. Therefore, we investigated whether presentation of monomer-delivered ligand was dependent on endosomal acidification. We pre-incubated BEAS-2B WT cells with bafilomycin before performing the exSPOT. Bafilomycin inhibits vacuolar ATPases^30^ and diminishes the presentation of an endosomal processing-dependent 5-A-RU pro-drug to MR1T cells^12^. Whereas the bafilomycin concentrations we used here reduced the MR1T cell response to this pro-drug in a dose-dependent manner, presentation of the monomer-delivered 5-OP-RU was not affected by the inhibitor (Figure 6b). Together, these data support a model in which the transfer of 5-OP-RU delivered in the context of a soluble MR1 monomer onto the MR1 expressed by the APC can displace pre-bound 6-FP and is independent of endosomal acidification.

Transfer of monomer-bound 5-OP-RU requires breaking a covalent bond. Since acidification was not needed for ligand transfer (Figure 6b), we hypothesized that this process is catalyzed by an exchange chaperone akin to HLA-DM for MHC II^4,31^. TAPBPR, tapasin, and the adapter protein AP2A1 have been shown to bind MR1 and modulate its antigen presentation^7,32,33^. However, knock down (KD) of these candidate chaperones alone or together did not affect how efficiently monomer-delivered 5-OP-RU was presented to our MR1T cell clone (Supplementary Figure 6).

Vesicular trafficking between the cell surface and the endosomal compartments is mediated by soluble N-ethylmaleimide-sensitive factor attachment protein receptors (SNAREs)^34^. The SNARE syntaxin-4 (encoded by *STX4*) is required for exocytosis in various cellular contexts^35–38^ including the fusion of recycling endosomes to the plasma membrane in cytotoxic T cells^35^ and post-synaptic membranes on dendritic spines^36^. Our lab has previously shown that small interfering RNA-mediated knock down of *STX4* reduces MR1-mediated antigen presentation of *M. smegmatis* supernatant but not live *Mycobacterium tuberculosis*^10^. To investigate the involvement of this mediator of vesicular trafficking in the exSPOT, we generated BEAS-2B cells deficient for *STX4* using CRISPR/Cas9 technology. Consistent with previous results in the knock down system^10^, *STX4* KO cells presented *M. smegmatis* supernatant less efficiently than their Cas9-expressing parent cell line (Figure 6c, p<0.0001). Similarly, presentation of monomer-delivered antigen was reduced in the absence of the SNARE (Figure 6c, p=0.0032). These results implicate syntaxin-4 in the mechanism underlying the exSPOT and suggest that it may share mechanistic parallels with presentation of microbial ligand.

## Discussion

Distinct and complementary cellular pathways have been proposed for MR1 loading and trafficking^11,14,39,40^. While most research efforts have been focused on the ER pathway, increasing evidence supports exchange in a post-ER compartment as a mechanism by which MR1 acquires antigenic ligand(s). For example, both Karamooz et al. and McWilliam et al. have shown that MR1 molecules in cells pre-incubated with the MAIT cell antagonist 6-FP can present activating MR1 ligands upon subsequent incubation with agonistic antigen^6,10^. Pre-treatment with 6-FP counters the ability of BFA to block surface expression of MR1/5-OP-RU complexes, indicating re-loading in a post-ER compartment^6^. Indeed, pre-incubation with 6-FP enhances MR1-mediated presentation of bacterial supernatant, presumably by increasing the pool of MR1 molecules available for re-loading^10^. Using a fluorescence internalization probe, McWilliam et al. also directly demonstrated recycling of a small proportion of MR1 molecules^6^. Furthermore, a recently developed pro-drug requires endosomal processing to release the 5-OP-RU precursor 5-A-RU^12^. Since the antigen is generated in endosomes in this scenario, it appears likely that the ligand is loaded onto MR1 in this compartment as opposed to trafficking to another intracellular organelle, which would require it to cross multiple membranes. Interestingly, while Ac-6-FP inhibits T cell activation after delivery of free 5-A-RU, it does not interfere with the presentation of this pro-drug^12^. This is consistent with the earlier observation that 6-FP blocks presentation of bacterial supernatant but not of intracellular infection^9^. The same study demonstrated that MR1- mediated antigen presentation of *M. smegmatis* supernatant is reduced upon knock-down of endosomal trafficking molecules^9,10^, further supporting involvement of this pathway in efficient loading of some antigens. In addition, a synthetic MR1 ligand modified to be able to track its location by microscopy resides in small vesicles also occupied by the micropinocytosis marker dextran^7^. This supports the idea that both membrane-bound internalized MR1 and soluble MR1 ligands could occupy this compartment at the same time, facilitating loading^14^. Together, these data lead us to hypothesize that there are distinct pools of MR1 in the ER and in a post-ER compartment^11,12,14,40^, that these compartments are differentially accessed by exogenous ligands, and/or that conditions are more favorable for ligand exchange in the endosomal compartment than in the ER. This notion is not without precedent as multiple, complementary pathways including both direct loading in the ER and ligand exchange in the endosomal compartment have been shown for MHC I (reviewed e.g. in ^41^).

In this study, we report a delivery method for MR1 ligands which allowed us to directly demonstrate transfer of the MR1T cell-activating ligand 5-OP-RU from recombinant, soluble MR1 onto cellular MR1. Intriguingly, monomer-delivered antigen was presented more efficiently than tetramer-delivered ligand. We speculate that the bulky structure of the tetramer may result in steric hindrance of interactions with chaperones required for ligand transfer. The smaller, more compact monomer, on the other hand, may be better suited to adopt the correct orientation. Moreover, DCs were much more efficient at presenting the tetramer-derived ligand than other cell lines tested. This could be due to their ability to cross-present internalized antigen, which allows their MHC I molecules to sample diverse sources of antigen and may enable MR1 to do likewise^18^.

Monomer-delivered 5-OP-RU was also presented to MR1T cells more efficiently than free 5-OP-RU. One explanation for this observation is that binding to the monomer protects the inherently unstable antigenic molecule from dehydrative cyclization^8^. However, the analogous results with MHC II tetramers argue that the enhanced antigen presentation associated with monomer delivery is not simply attributable to increased stability as peptide antigens are more stable. Therefore, we think it more likely that the monomer delivery enhances the efficiency of presentation by improving another aspect of antigen processing or presentation.

Previous 6-FP pre-loading experiments suggested that MR1 ligand exchange requires internalization of the antigen presentation molecules as it did not occur if cells were incubated at 4°C^6^. Interestingly, recent data from a similar model indicated that internalization mediated by the adapter protein AP2A1 is not required for the exchange of 6-FP for an MR1 ligand linked to a fluorophore^33^. While ligand exchange may occur on the cell surface in this model, the data presented in this manuscript support the conclusion that internalization of the ligand-loaded monomer is required for the exSPOT (Figure 4a-c; Figure 6a). Thus, it appears unlikely that the ligand exchange in this delivery method could occur at the plasma membrane.

Our efforts to identify the molecular mechanism that enables transfer of the antigenic ligand from the soluble MR1 to the cellular MR1 show that this process is dependent on actin dynamics and enhanced if cellular MR1 is pre-loaded with 6-FP. Peptide editing on MHC I and MHC II depends on the relative affinities of the available ligands and their competition for transiently ligand-receptive antigen presenting molecules^3,4,31^. In line with this concept, we propose that the endosomal compartment is a uniquely suited place for MR1 ligand exchange by virtue of its high local concentration of antigens derived from internalized pathogens relative to the cell as a whole. This enrichment of otherwise low abundance ligands may drive the equilibrium from pre-bound ligands to internalized antigens specifically in endosomal compartments.

This analogy breaks down, however, in that peptides are not covalently bound to their antigen presenting molecules. By contrast, a covalent bond needs to be broken in our exchange assay to release the antigenic ligand from the soluble MR1. The Schiff base that anchors both 6-FP and 5-OP-RU to the MR1 binding groove is considered labile under acidic conditions^6,14^. Nevertheless, our results suggest that ligand exchange on MR1 does not require endosomal acidification. This is in striking contrast to MR1-mediated presentation of fixed whole *E. coli*^13^ and the 5-A-RU pro-drug^12^. While the latter was designed to require acidic conditions for release, the first suggests that the molecular mechanisms for MR1 loading differ between *E.coli* and 5-OP-RU-loaded monomer. Importantly, *Mycobacterium tuberculosis* is known to prevent endosomal acidification to evade immune recognition^42^, making antigen loading mechanisms that function at higher pH desirable. Alternatively, ligand transfer after monomer delivery may be insensitive to bafilomycin because it occurs in the ER. Previous studies have demonstrated that whole proteins from the extracellular environment can be imported into this compartment^43–45^, so this may be true for the MR1 monomer, too. In that case, we would, however, expect monomer delivery to efficiently supply ligand to MHC I molecules as well, which is not the case (Figure 5c). Our BFA experiments also show that MR1 ligand exchange can take place after MR1 is released from the ER bound to 6-FP (Figure 5g).

While the molecular mechanism and the subcellular location of ligand exchange remain to be determined, our data lead us to propose that it is a facilitated process catalyzed by a yet unidentified exchange chaperone (Supplementary Figure 7). Monomer delivery may increase the efficiency of exchange by enhancing the interaction between pre-loaded MR1 and this chaperone. Indeed, specific chaperones facilitate the release of pre-loaded ligands, stabilize the transiently unoccupied antigen presenting molecule, and mediate the re-loading with antigenic cargo in other examples of ligand exchange. For MHC I, for instance, this function can be fulfilled by TAPBPR in the Golgi (reviewed e.g. in ^46^ and ^47^). For MHC II, HLA-DM chaperones endosomal ligand exchange and exchange kinetics are further modulated by its regulator HLA-DO (recently reviewed in ^31^ and ^4^). Recent screens for modulators of MR1 loading and trafficking have identified numerous interaction partners through both genetic techniques and protein co-immunoprecipitation^7,48^. These studies found MR1-mediated antigen presentation to be regulated by tapasin and TAPBPR^7,32^ and the P5-type ATPase ATP13A1^48^, but have largely focused on ER and Golgi-resident proteins. Discovery approaches more specifically targeted at interaction partners during ligand exchange in endosomal compartments may be needed to identify the chaperone(s) involved in this process.

In this manuscript, we explored the mechanisms by which MR1 ligands can be delivered exogenously using MR1 itself to stabilize an inherently unstable ligand. This mode of ligand delivery is a powerful tool to identify chaperones involved in the re-loading of pre-loaded MR1 molecules which can then be tested for their involvement in physiological settings such as mycobacterial infection. 5-OP-RU was also recently successfully used to boost mucosal immunity in the context of vaccinations against respiratory pathogens^49^. Thus, in addition to its use in exploring the molecular underpinnings of ligand exchange, we also postulate MR1/ligand complexes could be used as an adjuvant or antigen delivery strategy in vaccination approaches.

## Methods

### Human subjects

This study was conducted according to the principles expressed in the Declaration of Helsinki. Study participants, protocols, and consent forms were approved by the Institutional Review Board at Oregon Health & Science University (OHSU) (IRB00000186). All ethical regulations relevant to human research participants were followed. Peripheral blood mononuclear cells (PBMC) from human subjects were only used to make dendritic cells to be used as antigen presenting cells or used to expand T cell clones as described in the methods section. All findings in this manuscript are based on the T cell clone function or on other cell line function, e.g. BEAS-2B cell line. No data is presented on the PBMC from human subject participants and sex and gender were not taken into consideration.

### Cells and bacteria

*M. smegmatis* (strain mc^2^155) supernatant was obtained by growing *M. smegmatis*, centrifuging the cultures and filtering the supernatant through a syringe-driven 0.22 μm filter^10^. PBMCs were isolated from whole blood obtained by leukapheresis with informed consent. Human monocyte-derived dendritic cells (DCs) were isolated from PBMCs by plate adherence. For this, PBMCs were incubated in RPMI supplemented with 2% heat-inactivated human serum, 3.5 mM L-Glutamine (gibco), and 44 μg/ml gentamycin (gibco) at 37°C for 1h, before gentle tapping and removal of non-adherent cells. Adherent cell were then cultured in RPMI supplemented with 10% heat-inactivated human serum and L-Glutamine and gentamycin as above (RPMI10%HuS) and 30 ng/ml GM-CSF (Immunex) and 10 ng/ml IL-4 (R&D Systems) for 5 days^50^. BEAS-2B cells were obtained from the American Type Culture Collection (ATCC) and cultured as recommended. These cell lines are passaged up to 15 times before being discarded. Freezebacks are frozen down during the first 2-3 passages and are routinely tested for mycoplasma. BEAS-2B *MR1^-/-^*, BEAS-2B *MR1^-/-^*: *MR1A*^20,21^ and the BEAS-2B cell lines overexpressing GFP-tagged MR1A constitutively (BEAS-2B.MR1-GFP)^51^ or under the tetracycline-inducible promoter^52^ were described previously. THP-1 cells were obtained from ATCC. THP-1 *MR1^-/-^* and THP-1 *MR1^-/-^* : *MR1* cell lines were described previously^21,53^. The THP-1 *MR1^-/-^* from the Sewell lab constitutively express Cas9 protein whereas the THP-1 *MR1^-/-^* : *MR1* cells from the Cerundolo lab do not. EBV-transformed B cell lines (LCL) were generated in our laboratory using supernatants from the cell line 9B5–8 (ATCC). A549 cells were obtained from ATCC. The pCI AscI MR1 R9H res construct (Addgene ID 214753) was based on work by Howson et al.^24^. pCI AscI MR1 res (Addgene ID 214752) and pCI AscI MR1 R9H res (Addgene ID 214753) contain silent mutations to make the sequence resistant to CRISPR/Cas9 editing in the *MR1^-/-^* background. For both plasmids, the coding sequence with a downstream internal ribosomal entry site (IRES) GFP^54^ was ordered from GeneArt (ThermoFisher) and cloned into our previously described pCI AscI vector^10^. Plasmids were sequenced at the OHSU sequencing core and transfected into BEAS-2B *MR1^-/-^* cells using the Amaxa nucleofection system with Kit T solution (Lonza). The transfection efficiency was evaluated via GFP expression by flow cytometry (Supplementary Figure 4). *Trichoplusia ni* (Hi-5, Thermo Fisher Scientific) and *Spodoptera fugiperda* (Sf9, Thermo Fisher Scientific) cells for baculovirus production and recombinant protein expression were maintained as described previously^55^.

### CRISPR/Cas9-mediated knock out of STX4

To generate knockout cells, we used a two-plasmid system in which both the Cas9 gene and the sgRNA were cloned into different lentiviral vectors with independent selection markers similar to the method used by Niekamp et al. ^56^. The plasmids used for the generation of lentivirus comprise the lentiviral vector (pCDH-sgRNA or pCW-Cas9), the psPAX2 (packaging), and the pMD2.G (envelope) plasmids. The pCW-Cas9 plasmid was obtained from Addgene (#50661), as were psPAX2 (Addgene #12260) and pMD2.G (Addgene #12259). Lentiviruses were generated by transfecting low-passage HEK293T cells with these plasmids. Early passage BEAS-2B WT cells were stably transduced with these lentiviral particles encoding Cas9 by spinoculation^57^. Transduced cells were selected by culturing with 5 μg/ml Blasticidin (Life Tech) for 12 days. Three sgRNA constructs targeting *STX4* were generated using the sgOpti plasmid (Addgene ID 85681) as a backbone and packaged into lentiviral particles^58^. Early passage BEAS-2B.Cas9 cells were then transduced with supernatant containing these lentiviral particles in the presence of 200 μg Polybrene (Sigma). Transduced cells were selected by culturing with 8 μg/ml Puromycin (Sigma) for 6 days. Monoclonal populations were created by limiting dilution and a functional screen was performed by ELISPOT. Genomic editing efficiency and clonality were verified by Sanger sequencing. Briefly, DNA was isolated from *STX4* KO clones and BEAS-2B.Cas9 control cells using the QIAamp DNA Micro Kit (Qiagen) per manufacturer’s protocols. DNA surrounding the sgRNA target site was amplified by PCR (STX4-1F: ACAAGGTGGTTAAGGTGGCA; STX4-1R: CTGTTCACAGGGAGACCGAC) and purified using the QIAquick PCR Purification Kit (Qiagen) per manufacturer’s protocols. Sanger sequencing was performed by capillary electrophoresis at the OHSU Vollum Institute DNA Sequencing Core and sequences were analyzed by Tracking of Indels by DEcomposition (TIDE)^59^ and Inference of CRISPR Edits (ICE)^60^. Sequence of the sgRNA used to generate the *STX4* KO clonal cell line used in this manuscript: AGAACGTGGAGCGGATTCGG.

### T cell clones

T cell clones were expanded by co-culture with irradiated allogeneic PBMC and irradiated LCL in RPMI10%HuS in the presence of anti-CD3 antibody (Thermo Fisher Scientific, clone OKT3, Cat No. 16-0037-81, used at 30 ng/ml)^19,26^. Recombinant interleukin-2 (Proleukin, OHSU Inpatient Pharmacy) was added to the cultures every 2-3 days to a final concentration of 2 ng/ml and anti-CD3 antibody was washed off on Day 5. T cells were frozen down at -80°C at 1e6/ml after at least 11 days and used from frozen. T cell clones are expanded from frozen stocks using published protocols and are used for 2 weeks and then discarded. They are also tested to confirm their phenotype using either ELISPOT or flow cytometry and routinely tested for mycoplasma. Three T cell clones were used in this work. Two clones have been published previously (D426-G11^15,19^ and D480-F6^26,27^). D481-D1 is a CD4^+^ T cell clone that recognizes Esat6_73-87_. This clone was generated in a limiting dilution assay by adding either 100, 200 or 600 CD4^+^ T cells (CD4 microbeads, Miltenyi) isolated from PBMC from D481, a participant who had active tuberculosis, to wells of a 96 well plate containing 1e5 irradiated (3,500 rad using a Cs source) allogeneic PBMCs, 1e4 irradiated (3,500 rad using a Cs source) autologous dendritic cells (prepared as described above), 2 μg/ml Esat6 protein (BEI Resources), 5 ng/ml interleukin-2 and incubated at 37°C for two weeks. Wells with growth were assessed for response to an Esat6 peptide pool consisting of 15mers overlapping by 11 amino acids in an IFN-γ ELISPOT. T cell clones that responded to the Esat6 peptide pool were expanded as described above^19,26^ and further screened against the individual 15mers. The restricting allele is not known for this clone, but D481 expresses these class II alleles: DRB1*08:01, DRB1*13:01, DPB1*01:01, DPB1*04:01, DQB1*04:02 and DQB1*06:03 (HLA typing by Kashi Clinical Laboratories, Portland, OR).

### ELISPOT assays

For the plate-bound tetramer ELISPOT (tetraSPOT) assay, ELISPOT plates were coated with 50 μl of anti–IFN-γ antibody (Mabtech, clone 1-D1K, Cat No. 3420-3-1000, used at 10 μg/ml) per well. At the time of coating, MR1/5-OP-RU and MR1/6-FP tetramers conjugated to PE obtained from the NIH tetramer core^8^ were added at a range of concentrations. After overnight incubation at 4°C, ELISPOT plates were washed three times with phosphate-buffered saline (PBS) and then blocked with RPMI10%HuS for at least 1 h. T cell clones (1e3-2e4) were added to wells overnight. IFN-γ ELISPOTs were enumerated following development with an alkaline phosphatase-conjugated secondary antibody (Mabtech, clone 7-B6-1-ALP, Cat No. 3420-9A-1000, used 1:1000). Briefly, ELISPOT plates were washed extensively in PBS+0.05% Tween-20 (Thermo Scientific), incubated with secondary antibody for at least 2h at room temperature, washed in PBS+0.05% Tween-20 again and then PBS before incubation with BCIP/NBT-plus developer (Mabtech). Dried plates were imaged with an AID ELISPOT reader and spots were counted using the AID ELISPOT software Version 7.0. For all other ELISPOT assays, ELISPOT plates were coated with anti-IFN-γ antibody and blocked as described above. 1e4 DC, LCL, THP-1 or BEAS-2B cells were plated in the ELISPOT wells in RPMI10%HuS human serum. Tetramer, monomer, ligand or peptide were added to the wells at concentrations indicated for each experiment. PHA was used as positive control. After at least 1 h, 1e3-2e4 T cell clones were added and the plate was incubated overnight at 37°C. Antibody blocking was performed using the anti-MR1 26.5 clone (OHSU antibody core) and an IgG2a isotype control (Biolegend, clone MOPC-173, Cat No. 400224) added at 5 μg/ml for 1 h prior to the addition of ligand. 6-FP pre-treatment assays were performed as follows^10^. BEAS-2B.MR1-GFP cells were plated in 6-well tissue culture plates and treated with 6-FP *vs* NaOH control over night. On the next day, the cells were harvested and used as APCs in an IFN-γ ELISPOT assay as described above. For experiments with Bafilomycin A1 (MedChemExpress), BEAS-2B cells were incubated with Bafilomycin A1 for 2 h at indicated concentrations prior to the addition of 100 pM MR1/5-OP-RU monomer or 50 nM 5-OP-RU prodrug (MedChemExpress) in an IFN-γ ELISPOT assay as described above.

### 5-OP-RU Ligand

5-(2-oxopropylideneamino)-6-d-ribitylaminouracil (5-OP-RU) was made from two sources of 5-amino-6-D-ribitylaminouracil (5-A-RU). For the first method, 5-nitro-6-d-ribitylaminouracil (5-N-RU) was prepared by the OHSU Medicinal Chemistry Core. 5-A-RU was freshly prepared from 5-N-RU following a previously described procedure^8^. In brief, 4.8 mg of 5-N-RU was dissolved in 400 μl of 50 mM MES, pH 5.8. 8-fold molar excess of sodium dithionite powder was added and incubated in the dark at 90°C for five minutes under argon. The solution was chilled in an ice water bath. To prepare 5-OP-RU, at least 15 molar equivalents of methylglyoxal was added to 5-A-RU in 100 mM HEPES, pH 7.2, 150 mM NaCl to give a stock solution at 20 mM. For the second method, 5-A-RU*HCl was prepared by the OHSU Medicinal Chemistry Core following a previously described procedure^61^. To prepare 5-OP-RU, equal volumes of circa 650 mM methylglyoxal (Sigma) and 32 mM 5-A-RU*HCl dissolved in water were combined immediately before addition to the assay to give a stock solution of 16 mM of 5-OP-RU. Mass spectrometry analysis confirmed near-complete conversion of 5-A-RU into 5-OP-RU under similar conditions (Supplementary Figure 8). For this, samples were directly injected into an AB Sciex TripleTOF 5600 quadrupole time of flight mass spectrometer using the built-in syringe pump through a Hamilton Model 1710 N 100 µl syringe. Samples passing through the syringe were ionized by a PicoView ion source by New Objective before passing through a Co-Ann technologies 11cm 20 µm inner diameter pulled glass emitter which tapered to a 10 µm inner diameter at the tip. A stock solution of 32 mM 5-A-RU was diluted 1:10 before injection into the mass spectrometer. 5-OP-RU was produced by combining a 5 µl stock solution of 32 mM of 5-A-RU with a 5 µl 1:10 dilution from a 6.5 M stock solution of methylglyoxal (Sigma). This 5-OP-RU was then diluted 1:10 for injection into the mass spectrometer. Samples were examined in negative ion mode with the source voltage set at -2600 V. MS1 and MS2 data was acquired using Analyst 1.8 in manual tuning mode, with a TOF mass range of 30 to 500 Da after 1 minute. MS2 data was collected in high sensitivity mode with a declustering potential of -40 and a collision energy of -30 for 5-A-RU and -38 for 5-OP-RU. Data was analyzed using PeakView 1.2.

### Flow Cytometry assays

The following reagents were obtained through the NIH Tetramer Core Facility: MR1/5-OP-RU tetramer and monomer and MR1/6-FP tetramer and monomer, HLA*B08:01/CFP10_3-11_ tetramer, HLA-DRB1*14:03/Esat6_73-87_ tetramer, HLA-DRB1*14:03/control tetramer. Identification of MAIT cells with MR1 tetramers was described in ^8^. The D426-G11 TCR tetramer was made as specified below. To confirm the specificity of the TCR tetramer as a staining reagent, BEAS-2B.MR1-GFP cells were incubated with the indicated concentrations of 5-OP-RU or 6-FP, or the equivalent volume of 0.01 M NaOH overnight. Cells were harvested with Trypsin and split into three wells of a U-bottom 96-well plate each for staining with TCR tetramer, anti-MR1 antibody 26.5 (BioLegend, Cat No. 361108, used at 0.4 μg/ml), or isotype control antibody (BioLegend, Cat No. 400222, used at 0.4 μg/ml) for at least 30 min on ice. BEAS-2B *MR1^-/-^*cells transfected with the pCI AscI MR1 res or pCI AscI MR1 R9H res plasmids were analyzed similarly. For ligand titration experiments, 5e4 BEAS-2B.MR1-GFP cells per well were incubated with the indicated concentrations of free 5-OP-RU or MR1/5-OP-RU monomer in a 24-well plate overnight. Cells were harvested with Trypsin, transferred to a 96-well U-bottom plate, washed, and stained with TCR tetramer for at least 30 min at 4° C. For time course experiments, 5e4 BEAS-2B.MR1-GFP cells per well were plated in 24-well plates. Free 5-OP-RU or MR1/5-OP-RU were added to the indicated final concentrations for the indicated amounts of time. Samples from all time points were harvested and stained at the same time as above. Free ligand and monomer titrations were performed in parallel in both sets of experiments. For the actin blockade, 5e4 BEAS-2B.MR1-GFP cells were plated in 24-well plates and left to adhere overnight. Cells were then treated with 10 μM Latrunculin B (Sigma) or the equivalent volume of DMSO (Sigma). After 1 h, MR1/5-OP-RU or MR1/6-FP monomer was added to a final concentration of 100 nM and the cells were incubated for another 4 h before staining as above. For Brefeldin A blocking experiments, BEAS-2B.MR1-GFP cells were incubated with 94 μM 6-formylpterin (6-FP, Schircks) or the equivalent volume of 0.01 M NaOH (Sigma) overnight at 37°C at 5e4/well in a 24-well plate. 6-FP was washed off and the cells treated with 50 μg/ml Brefeldin A (BFA, Sigma) or the equivalent volume of DMSO (Sigma) for 1 h, followed by the addition of MR1/6-FP or MR1/5-OP-RU monomer (100 nM) or free 5-OPRU (5 μM) or 6-FP (100 μM) and incubation at 37°C for 4 h. Cells were then stained with TCR tetramer as above. The gating strategy is shown in Supplementary Figure 9. TCR tetramer was used at 150 or 300 nM. BEAS-2B WT cells were used for unstained and single color controls. Anti-MHC I antibody W6/32 (BioLegend, Cat No. 311410, used at 0.64 or 2.56 μg/ml) was used for the APC single color control due to low MR1 expression on BEAS-2B WT cells. For surface staining of BEAS-2B.MR1-GFP cells with streptavidin, 3e5 cells were incubated with 50 nM of MR1/5-OP-RU monomer or medium overnight. These cells were split into multiple wells and either stained with the TCR tetramer as described above or blocked with PBS + goat, human, and bovine serum for at least 30 min on ice and then stained with Streptavidin-APC (Agilent) for 40 min on ice or with biotinylated anti-MR1 antibody 26.5 (BioLegend, custom conjugation, used at 5 μg/ml) for 40 minutes on ice, followed by Steptavidin-APC for 40 minutes on ice. For tetramer staining of the CD8 T cell clone D480-F6, 1e5 T cell clones were incubated with the HLA*B08:01/CFP10_3-11_ tetramer at 1:500 (0.127 μg/well) for 45 minutes at 37°C. For tetramer staining of the CD4 T cell clone D481-D1, 1e5 T cell clones were incubated with the HLA-DRB1*14:03/Esat6_73-87_ or HLA-DRB1*14:03/control at 6 μg/ml for 1 h at 37°C. Cells were subsequently stained with an antibody cocktail containing LIVE/DEAD Fixable Dead Cell Stain Kit (Thermo Fisher), and surface stained with the antibodies to CD4 (BD Biosciences, clone SK3, Cat No. 348789 or 612748, used at 2.5 or 1 μl/test, respectively), CD8 (Biolegend, clone SK1, Cat No. 344714, used at 0.01 μl/test), and CD3 (BD Biosciences, clone SK7, Cat No. 340663, used at 3 μl/test or BioLegend, clone OKT3, Cat No. 317324, used at 1 μl/test), for 30 minutes at 4°C. Samples were washed and fixed with 1% PFA. Acquisition was performed using a BD FACSymphony, LSRII, or Canto cytometer. The gating strategy is shown in Supplementary Figure 10. Flow cytometric analysis was performed using FlowJo version 10.6.1 or 10.7.1 (Treestar, Ashland, OR, USA). Plots show GFP+ gates to ensure only cells over-expressing MR1 are included in the analysis.

### Knock down of candidate chaperones

For small interfering RNA-mediated knock down of candidate chaperones, small interfering RNAs targeting TAPBPR (Assay ID s229969), tapasin (Assay ID 41850), or AP2A1 (Assay ID s183) or a missense control (4390844) (all Thermo Fisher Scientific) were nucleofected into 1e6 BEAS-2B WT cells at 300 nM according to the manufacturer’s instructions (Lonza, Nucleofector Kit T). Cells were allowed to rest overnight (TAPBPR and tapasin) or two days (AP2A1) before use in an ELISPOT assay as described above. RNA levels of targeted transcripts were measured by qRT PCR at the time of ELISPOT using Taqman probes Hs00794155_m1, Hs00917451_g1, and Hs00900330_m1 with GAPDH (Hs02758991_g1) as the housekeeping gene (all Thermo Fisher Scientific). For AP2A1, equal cell numbers were also lysed in Lysis Buffer (Chromotek) with Protease Inhibitor cocktail (Roche) at the time of ELISPOT and lysates of all three experiments were analyzed together by Western blot. Briefly, lysates were combined with Loading Buffer and Reducing Agent (both Invitrogen), separated on 4-20% polyacrylamide gels (Bio-Rad), and transferred onto PVDF membrane (Millipore). The membrane was blocked with Blocking Buffer (Odyssey) for 1 h at room temperature before incubation with mouse anti-AP2A1 (BD, clone 8, Cat No. 610501, used at 250 ng/ml) and rabbit anti-ACTB (Abcam, Cat No. ab8227, used at 200 ng/ml) primary antibodies overnight at 4° C. The blot was developed with LI-COR IRDye secondary antibodies (goat anti-mouse 800CW and goat anti-rabbit 680RD, Cat No. 926-32210 and 926-68071, both used at 100 ng/ml) and imaged simultaneously on an Odyssey imager.

### Recombinant protein expression and purification

All experiments using MR1 monomer and tetramer were performed with the reagent(s) provided by the NIH Tetramer Core Facility. The D426-G11 TCR was produced as previously described for other MAIT TCRs^62^. Briefly, the D426-G11 TCR V(D)J region of each TCR chain was cloned separately into the pAcGP67a vector after the GP67 signal sequence and before either a “cysteine-optimized” TRAC or TRBC sequence^63^. To ensure heterodimer formation, the chains were tagged with cleavable C-terminal acidic and basic zippers^64^ each followed with a 6xHis tag. The alpha chain construct contained a C-terminal AviTag before the zipper cleavage site. The P1 and P2 baculoviruses were produced in Sf9 cells^55^, then the baculoviruses were added in tandem to Hi-5 cells and the cells were allowed to express the recombinant protein for 72 h. The supernatant was harvested and the protein purified by Ni-NTA chromatography. Recombinant TCR was then treated with recombinant HRV 3C protease overnight and the cleaved zippers were removed with Ni-NTA. The protein was biotinylated *in vitro* using recombinant BirA ligase and the biotinylation efficiency checked by a streptavidin gel-shift assay. Finally, it was purified by size exclusion chromatography and tetramerized with streptavidin-APC (Agilent). This reagent is available upon request after the completion of an MTA that includes a clear description of how the material would be used.

### Microscopy

BEAS-2B cells (1e5) were seeded on a chamber slide and incubated for 18 h at 37C. MR1/5-OP-RU tetramer conjugated to AF647 (200 nM) was added to the experimental wells for 2 h at RT. Control wells received no tetramer. To determine whether the tetramer was observed in the cells, CellMask Orange Plasma (1:5000 dilution, Invitrogen) was added for 10 min (Supplementary Figure 11). To examine whether the tetramer localized to endosomal compartments, CellLight BacMam 2.0 reagents (Molecular Probes) for early endosomes (Rab5 red fluorescent protein (RFP), Cat No. C10587) or lysosomes (LAMP1 RFP, Cat No. C10597) were added for 18 h prior to addition of the tetramer. For all imaging, DAPI was added during PBS washes, and cells were fixed prior to data acquisition. Images were acquired on a Keyence BZ-X810 Fluorescence microscope or high-resolution wide-field CoreDV microscope (Applied Precision) with CoolSNAP ES2 HQ (Nikon), using nuclear staining with DAPI to identify un-biased fields of view. Localization of the tetramer to endosomal compartments was quantified using Imaris image analysis software ^50^.

### Data analysis, Statistics, and Reproducibility

Data were analyzed with Prism 9 GraphPad Software. For measurements of functional avidity, curves were determined by non-linear regression and best-fit EC_50_ was calculated. EC_50_ values were compared with the extra sum-of-squares F test. Number of experimental and technical replicates are specified in the figure legends.

### Data Availability

The authors declare that the data supporting the findings of this study are available within the paper and its supplementary information files. Plasmids are available via Addgene with the ID numbers provided in the Methods section.

## Acknowledgements

We would like to thank the participants who gave time and dedication to this health research as well as Erin Merrifield, Department of Pediatrics, OHSU, for her contributions to this study.

This project has been funded in whole or in part with Federal funds from the National Institutes of Allergy and Infectious Diseases, National Institutes of Health, Department of Health and Human Services, under grant no AI147954 (EA, DML, WHH), AI134790 (DML), GM007183 (NAL), AI129976 (MH), R01AI141549-01A1 (FT) and AI165618 (AO). This work was also supported in part by Merit Award #I01 CX001562 from the U.S. Department of Veterans Affairs Clinical Sciences Research and Development Program (MJH), Merit Award #I01 BX000533 from the U.S. Department of Veterans Affairs Biomedical Laboratory (DML).

## Author Contributions

CK, GS, NAL, DAL, EJA, DML and MH contributed to the conception and/or design of the work. CK and GS are joint first authors. DML and MH jointly supervised the work. MC, MN, DAL and DML contributed to clinical activities. AO, WHH, FT, EJA, DML and MH raised grants to fund the research. CK, GS, NAL, MC, MN, AW, CL, TA, JB, TNL, CMH, MEH, JK, LA, AO, WHH, SM, FT, DAL, EJA, DML and MH substantially contributed to the acquisition, analysis, or interpretation of data and drafting of the manuscript. All authors substantially contributed to revising and critically reviewing the manuscript for important intellectual content. All authors approved the final version of this manuscript to be published and agree to be accountable for all aspects of the work.

