## Supplemental Information for "Delivery of loaded MR1 monomer results in efficient ligand exchange to host MR1 and subsequent MR1T cell activation"

### Affiliations

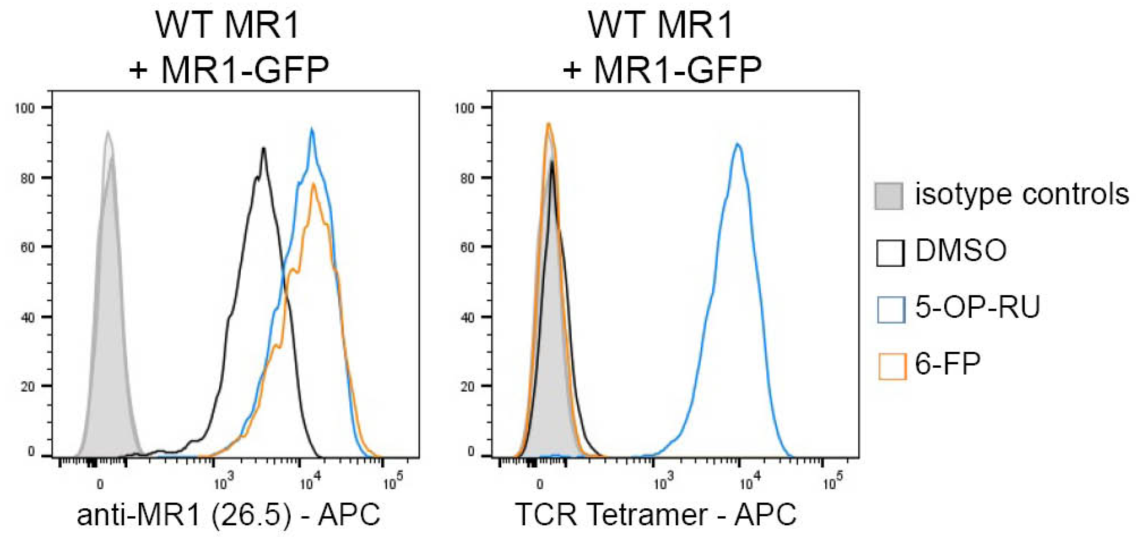

**Supplementary Figure 1. The TCR tetramer stains cells presenting 5-OP-RU but not 6-FP.**
BEAS-2B.MR1-GFP cells were incubated with 10.5 µM 5-OP-RU, 100 µM 6-FP, or the equivalent volume of 0.01 M NaOH overnight in 6-well plates. Cells were then harvested and surface stained with APC-conjugated TCR tetramer or APC-conjugated anti-MR1 antibody. Data in Supplementary Fig. 1 and 3 were collected together and are representative of n=3 independent experiments. The same isotype control samples are shown for TCR and antibody staining.

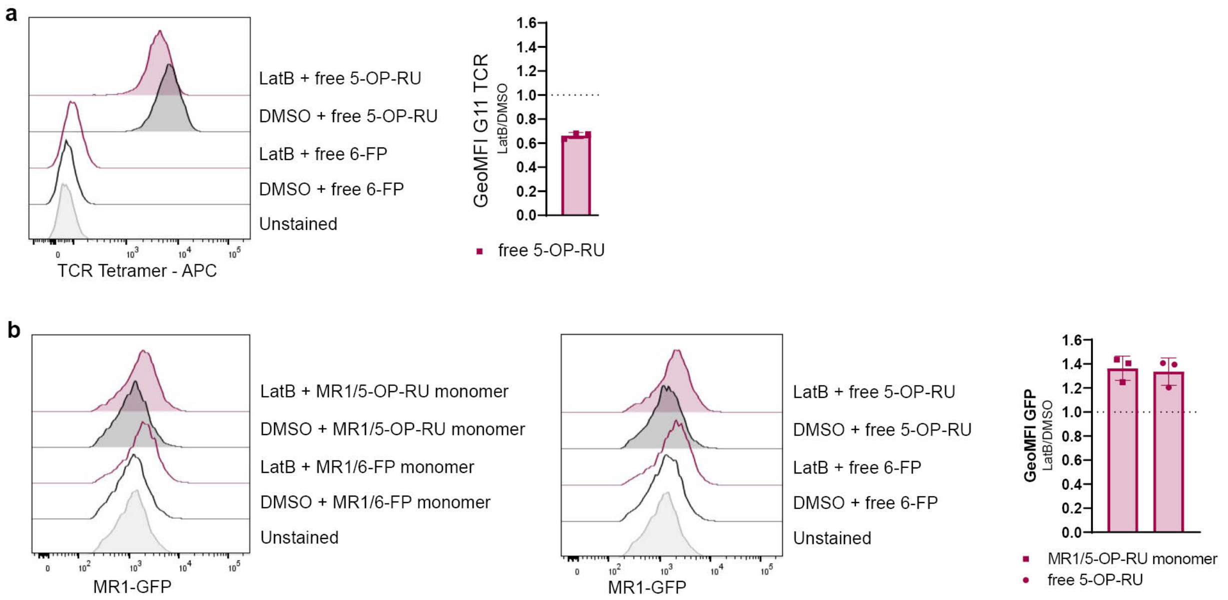

**Supplementary Figure 2. LatB treatment also impedes presentation of free 5-OP-RU but does not reduce total MR1 expression.**

The experiments shown in Figure 4b also included a free 5-OP-RU condition. **a.** BEAS-2B.MR1-GFP cells were pre-incubated with 10 µM of latrunculin B (LatB) for 1 h before addition of 5-OP-RU or 6-FP for an additional 4 h. Cells were surface stained with TCR tetramer. **b.** GFP data for the same cells and those in Figure 4b. Representative histograms are shown with pooled geometric mean fluorescence intensity (GeoMFI) from n=3 experiments. Error bars denote SD. Bars denote mean. The same unstained control is shown in all plots (including Figure 4b).

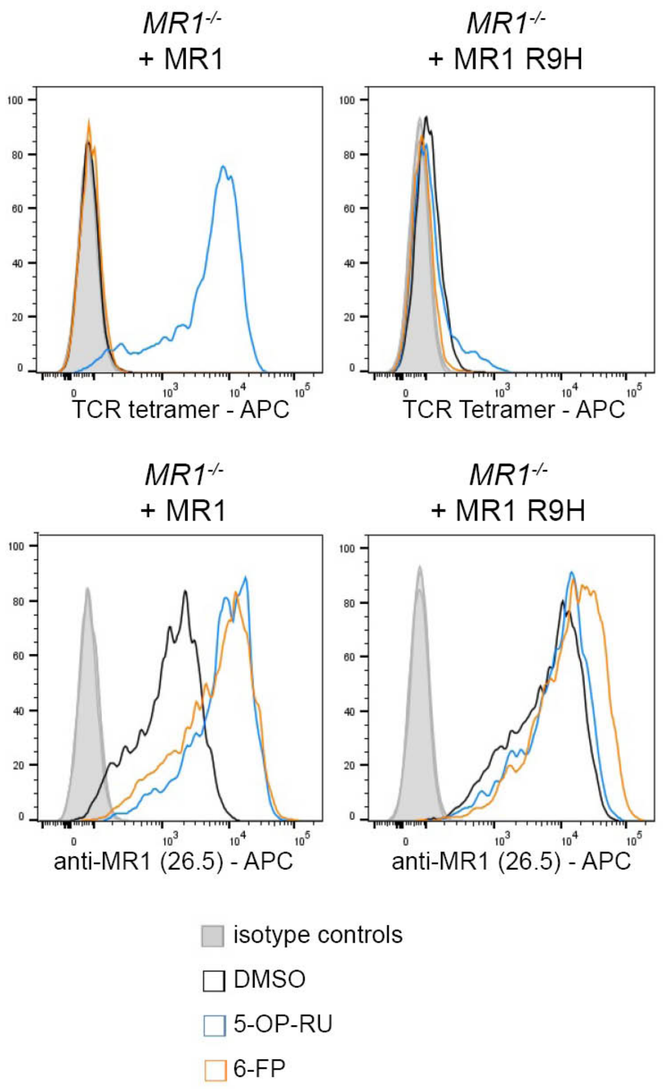

**Supplementary Figure 3. BEAS-2B *MR1*$^{-/-}$ cells transfected with MR1 R9H are unable to present 5-OP-RU.**

BEAS-2B *MR1*$^{-/-}$ cells were transfected with the constructs encoding MR1 or MR1 R9H upstream of an IRES GFP sequence as in Fig. 4. 1e5 cells were incubated with 10.5 µM 5-OP-RU, 100 µM 6-FP, or the equivalent volume of 0.01 M NaOH overnight in 6-well plates. Cells were then harvested and surface stained with APC-conjugated TCR tetramer or APC-conjugated anti-MR1 antibody. Data in Supplementary Fig. 1 and 3 were collected together and are representative of n=3 independent experiments. The same isotype control samples are shown for TCR and antibody staining.

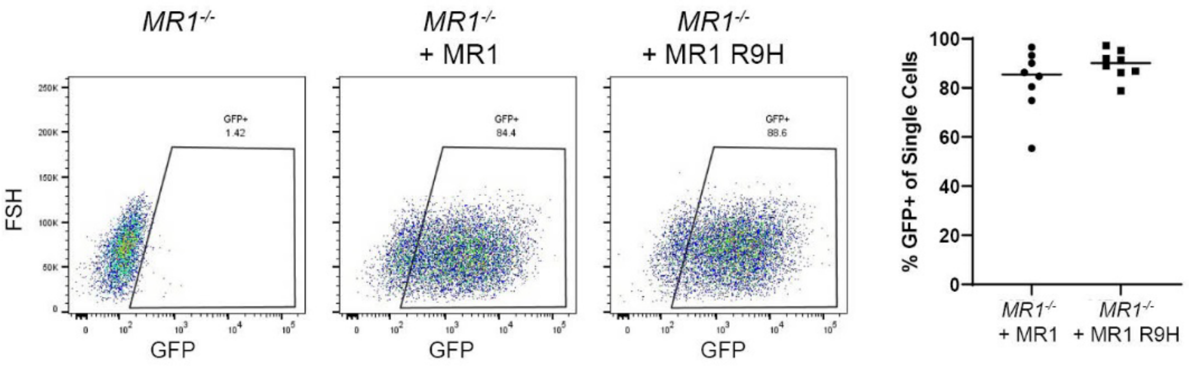

**Supplementary Figure 4. Transfection efficiencies are not lower for MR1 R9H than MR1.**
BEAS-2B *MR1*$^{-/-}$ cells were transfected with the constructs encoding MR1 or MR1 R9H upstream of an IRES GFP sequence as in Fig. 4 and analyzed for GFP expression by flow cytometry. A representative dot plot with gating is shown with percentages pooled from n=8 different experiments including those shown in Fig. 4 and Supplementary Fig. 2. Bars denote median. Cell number and additional staining differ between experiments. FSH = forward scatter height.

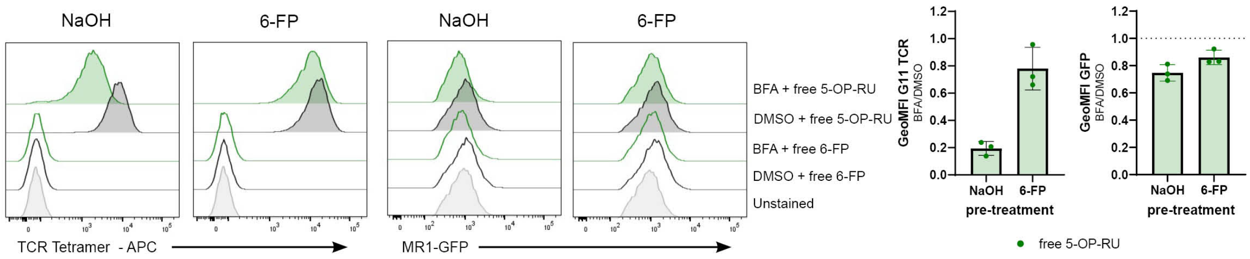

**Supplementary Figure 5. BFA treatment also impedes presentation of free 5-OP-RU and reduces total MR1 expression.**

The experiments shown in Figure 5g also included a free 5-OP-RU condition. BEAS-2B.MR1-GFP cells were pre-incubated with 6-FP or NaOH overnight. 6-FP was washed off and cells were incubated with 50 ug/ml brefeldin A (BFA) for 1h before addition of free 5-OP-RU or 6-FP for an additional 4 h. Cells were surface stained for flow cytometry with TCR tetramer. Representative histograms are shown with pooled geometric mean fluorescence intensity (GeoMFI) from n=3 experiments. Error bars denote SD. Bars denote mean. The same unstained control is shown in all plots (including Figure 5g).

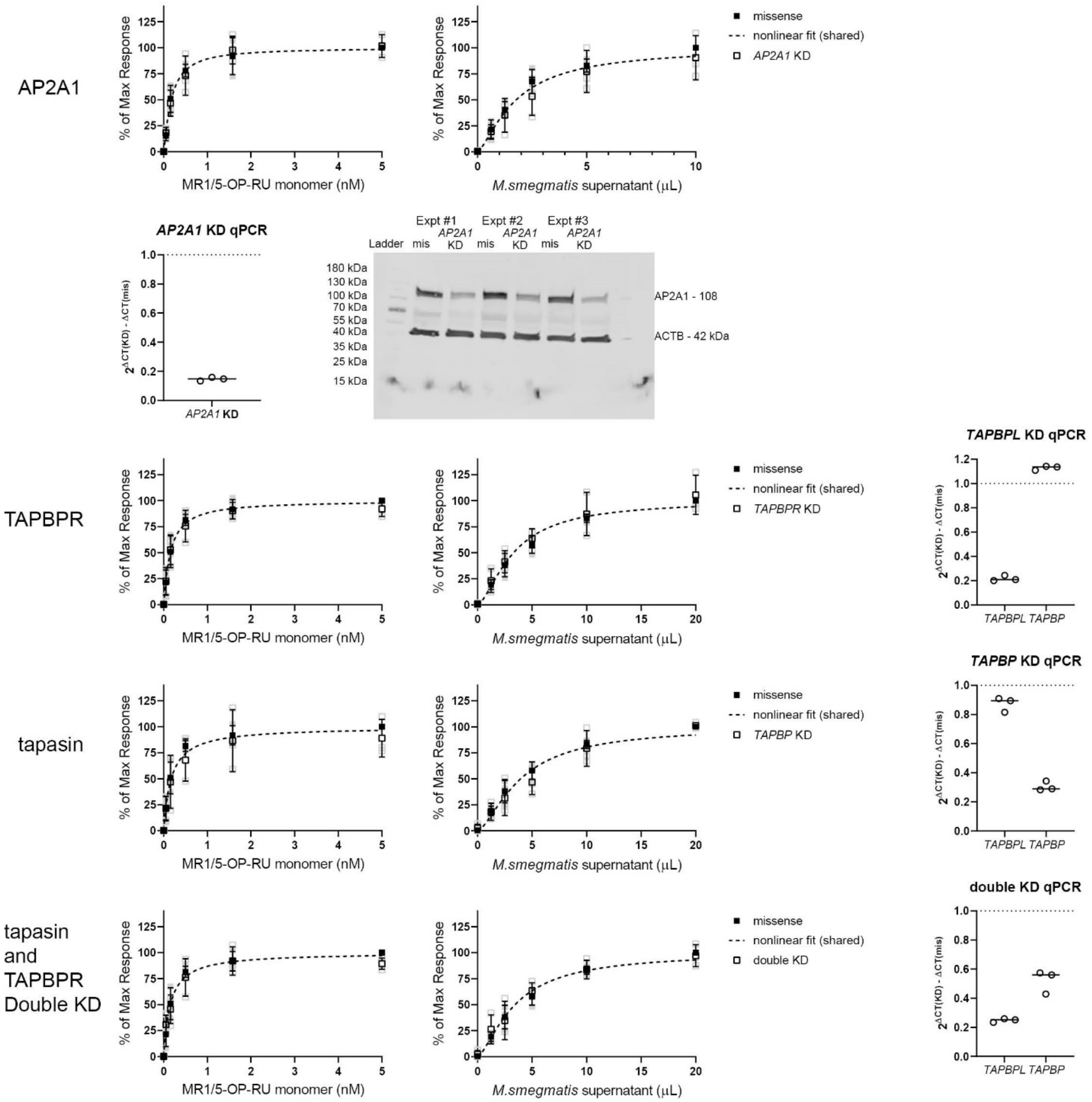

**Supplementary Figure 6. Knock down of *TAPBPL*, *TAPBP*, or *AP2A2* does not affect the exSPOT.** MR1T clone (1e3) IFN-$\gamma$ response to BEAS-2B WT cells transfected with siRNAs targeting the indicated mRNAs in the presence of the indicated concentrations of *M. smegmatis* supernatant or MR1/5-OP-RU monomer. Data were normalized to the maximum response and pooled from n=3 independent experiments. Error bars denote SD of the mean of pooled experiments which are shown individually in lighter colors. Non-linear regression analysis with extra sum-of-squares F test detected no statistically significant differences in nonlinear fit between missense and KD cell lines (*AP2A1*: p=0.1326 for supernatant, p=0.69 for monomer; *TAPBPL*: p=0.54 for supernatant, p=0.6372 for monomer; *TAPBP*: p=0.3046 for supernatant, p=0.3204 for monomer; double KD: p=0.8194 for supernatant; p=0.3394 for monomer). The same missense controls are shown for *TAPBPL, TAPBP*, and double KD as these experiments were performed together. Transcript levels at the time of plating for the ELISPOT were determined for each experiment and are shown with median. For *AP2A1* KD experiments, protein levels were also determined by Western Blot. IRDye-coupled secondary antibodies specific to the respective host species of the primary antibodies were used to image both proteins together but both are shown in black.

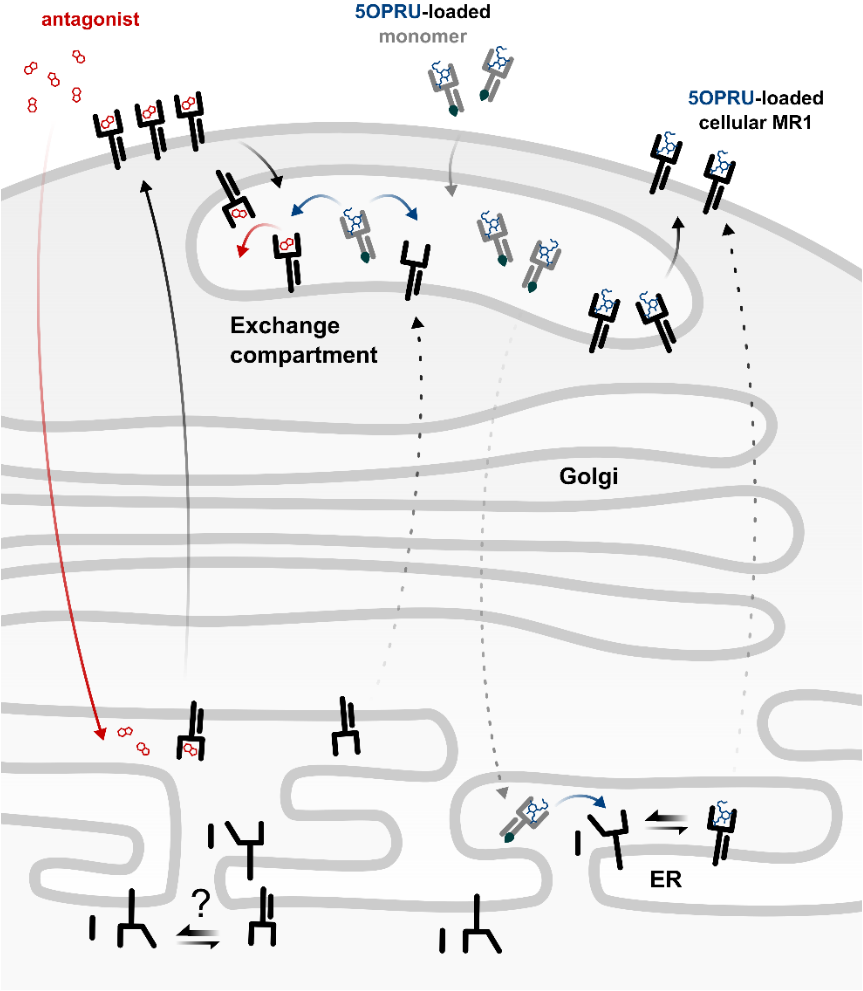

**Supplementary Figure 7. Conceptual model of the exSPOT.** We postulate that binding to a non-activating ligand (such as 6-FP in Figure 5a or an endogenous ligand at baseline) allows ER egress of loaded cellular MR1 molecules, which are then available for ligand exchange in a yet-to-be-defined Exchange compartment. 5-OP-RU-loaded monomer is trafficked to this compartment where the antagonist is exchanged for the antigenic ligand before the cellular MR1 returns to the cell surface for antigen presentation. The mechanisms of exchange do not involve acidification, TAPBPR, tapasin, or AP2A1 in our model but likely rely on an exchange chaperone that remains to be identified. Alternatively, the 5-OP-RU-loaded MR1 monomer may traffic to the ER through retrograde transport and the antigen could be loaded onto ligand-receptive MR1 molecules there before translocating to the cell surface.

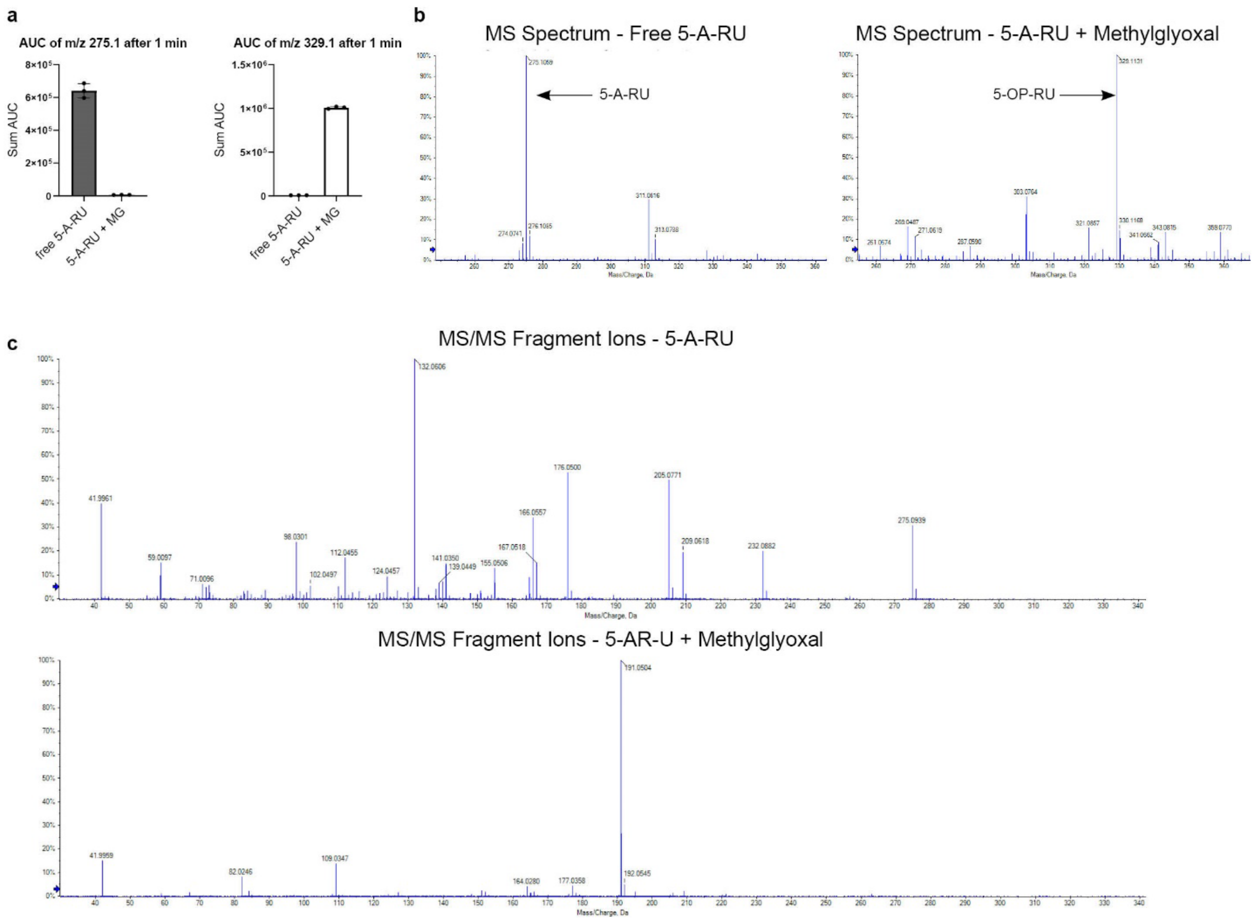

**Supplementary Figure 8. Monitoring conversion of 5-A-RU to 5-OP-RU by mass spectrometry.**
**a**. Area under the curve (AUC) analysis of direct infusion of 5-A-RU or 5-A-RU combined with methylglyoxal. Error bars denote SD of the mean of n=3 separate reactions. Bars denote mean. **b**. MS1 spectrum of 5-A-RU or 5-A-RU combined with methylglyoxal. **C**. MS2 fragment spectrum of 5-A-RU or 5-A-RU combined with methylglyoxal.

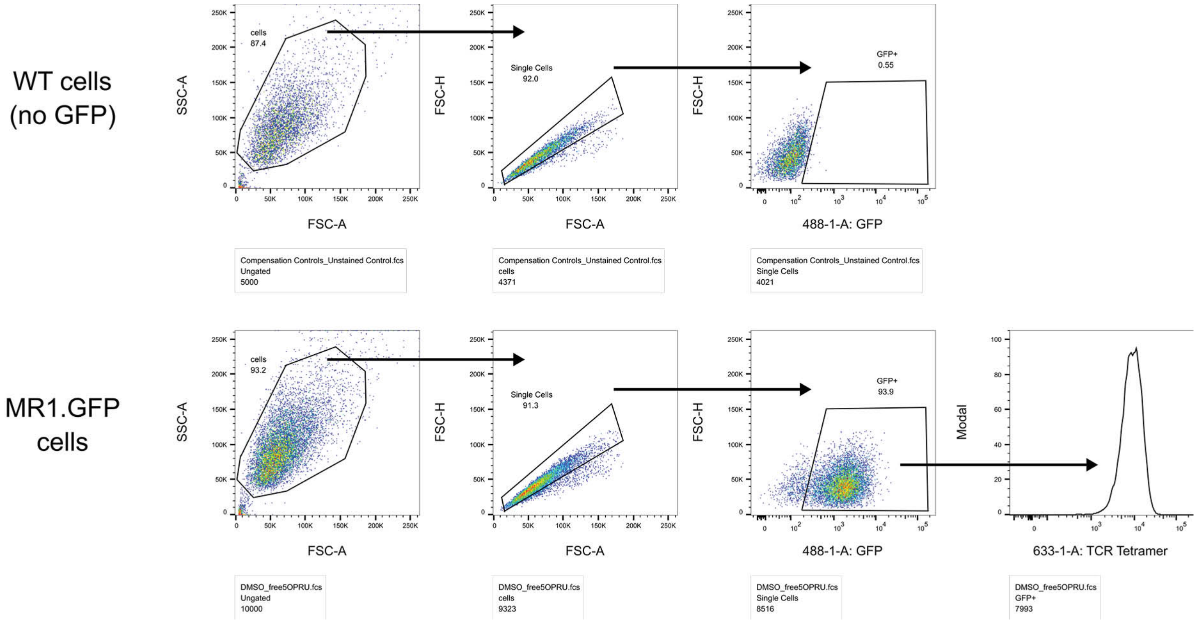

**Supplementary Figure 9. Gating strategy for BEAS-2B cells.** BEAS-2B.MR1-GFP cells were incubated with 5 µM 5-OP-RU for 4 h and surface stained for flow cytometry with APC-conjugated TCR tetramer as in Figure 4B. The same gating strategy was used for all flow cytometry experiments using BEAS-2B cells and TCR tetramer. SSC-A = side scatter area. FSC-A = forward scatter area. FSC-H = forward scatter height.

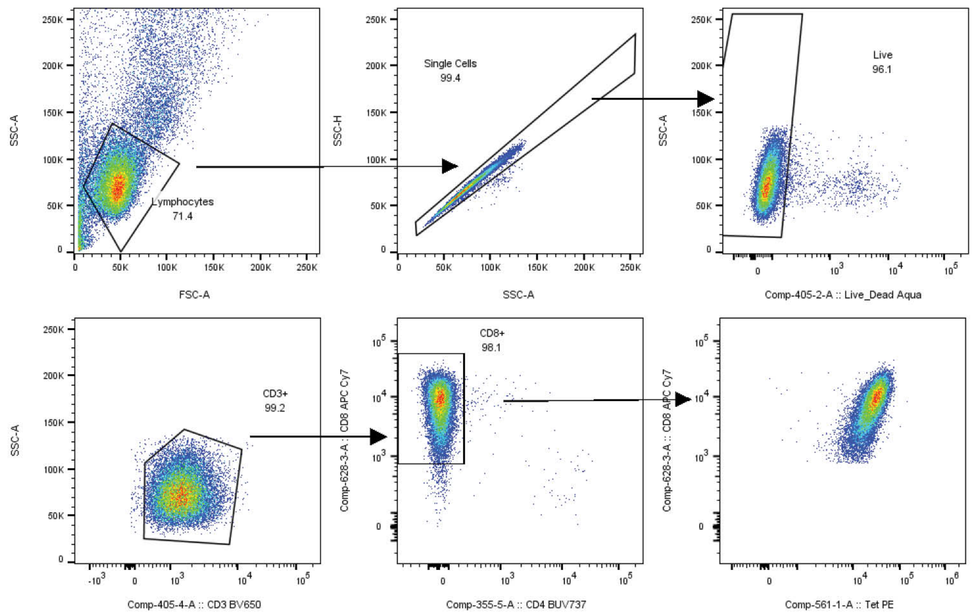

**Supplementary Figure 10. Gating strategy for CD8 or CD4 T cell clone.** HLA*B08:01-restricted T cell clone D480-F6 was stained with the HLA-B*08:01 tetramer to CFP10$_{3\text{-}11}$, along with antibodies to CD3, CD4, CD8 and a live/dead discriminator. The same gating strategy was used for all flow cytometry experiments with CD8 or CD4 T cell clones.

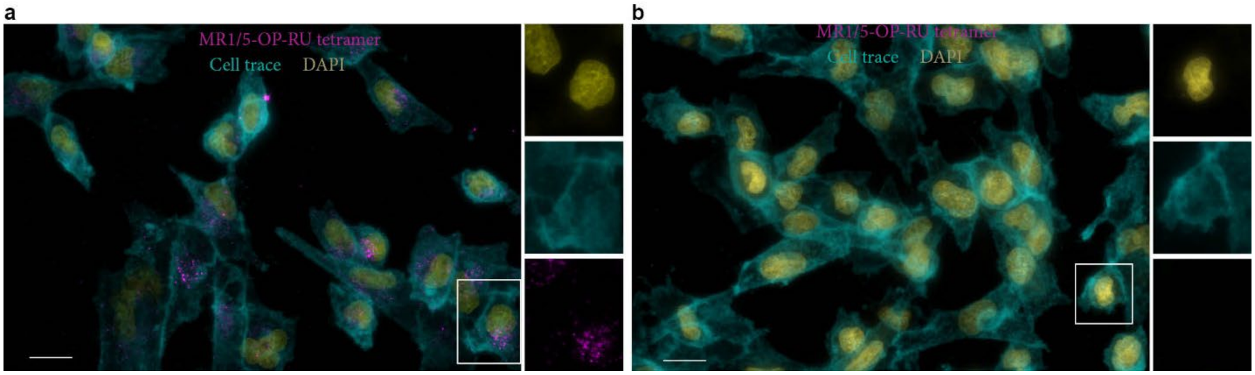

**Supplementary Figure 11. Specificity of Tetramer signal in Imaging.**
BEAS-2B cells (1e5) were seeded on chamber slide and incubated for 18 h at 37C. MR1/5-OP-RU tetramer conjugated to AlexaFluor647 (200 nM) was then added for 2 h at room temperature (**a**). Control wells received no tetramer (**b**). All cells then had CellMask Orange Plasma (1:5000 dilution, Invitrogen) added for 10 min before DAPI was added and cells were washed 2X in PBS. All images acquired were using the same microscope settings. Scale bar denotes 20 µm. Data are representative of n=3 independent experiments.

**Supplementary Table 1. EC50 values for Figure 1b.**

| | EC50 | | p-value |
|---|---|---|---|
| Expt | $\alpha$MR1 | IgG2a | $\alpha$MR1 vs IgG2a |
| 1 | 31.62 | 0.9504 | <0.0001 |
| 2 | 31.28 | 0.9319 | <0.0001 |
| 3 | 36.26 | 0.4913 | <0.0001 |
| Mean | 33.05 | 0.7912 | |
| Std Dev. | 2.782 | 0.2599 | |

**Supplementary Table 2. EC50 values for Figure 1c.**

| | EC50 | | | | p-value | | |
|---|---|---|---|---|---|---|---|
| Expt | Dendritic Cells | THP-1 | LCL | BEAS-2B | DC vs. THP-1 | DC vs. LCL | DC vs. BEAS-2B |
| 1 | 0.5550 | 141.7 | 255.0 | 264.8 | <0.0001 | <0.0001 | <0.0001 |
| 2 | 0.09276 | 43.26 | 124.6 | 348.2 | <0.0001 | <0.0001 | <0.0001 |
| 3 | 0.2645 | 154.3 | 115.0 | 192.8 | <0.0001 | <0.0001 | <0.0001 |
| Mean | 0.2790 | 113.1 | 164.9 | 268.6 | | | |
| Std Dev. | 0.1973 | 60.80 | 78.21 | 77.77 | | | |

**Supplementary Table 3. EC50 values for Figure 2a.**

| | EC50 | | | p-value | | |
|---|---|---|---|---|---|---|
| Expt | MR1/ 5-OP-RU tetramer | MR1/ 5-OP-RU monomer | 5-OP-RU | MR1/5-OP-RU tetramer vs. MR1/5-OP-RU monomer | MR1/5-OP-RU tetramer vs. 5-OP-RU | MR1/5-OP-RU monomer vs. 5-OP-RU |
| 1 | 0.2038 | 0.2164 | 4.967 | 0.7517 | <0.0001 | <0.0001 |
| 2 | 0.06801 | 0.06178 | 3.746 | 0.6100 | <0.0001 | <0.0001 |
| 3 | 0.07956 | 0.1002 | 5.654 | 0.3384 | <0.0001 | <0.0001 |
| Mean | 0.1171 | 0.1261 | 4.789 | | | |
| Std Dev. | 0.07529 | 0.08050 | 0.9664 | | | |

**Supplementary Table 4. EC50 values for Figure 5a.**

| | EC50 | | p-value |
|---|---|---|---|
| Expt | 6FP | NaOH | 6FP vs NaOH |
| 1 | 0.1365 | 1.027 | <0.0001 |
| 2 | 0.1372 | 0.8863 | <0.0001 |
| 3 | 0.2087 | 3.154 | <0.0001 |
| Mean | 0.1716 | 1.689 | |
| Std Dev. | 0.04012 | 1.271 | |
